# Early β-amyloid accumulation in the brain is associated with peripheral T cell alterations

**DOI:** 10.1101/2023.01.17.524355

**Authors:** Christoph Gericke, Tunahan Kirabali, Roman Flury, Anna Mallone, Chiara Rickenbach, Luka Kulic, Vinko Tosevski, Christoph Hock, Roger M. Nitsch, Valerie Treyer, Maria Teresa Ferretti, Anton Gietl

## Abstract

Fast and minimally invasive approaches for early, preclinical diagnosis of neurodegenerative Alzheimer’s disease (AD) are highly anticipated. Evidence of adaptive immune cells responding to cerebral β-amyloidosis, one of the pathological hallmarks of AD, has raised the question of whether immune markers could be used as proxies for β-amyloid accumulation in the brain. Here, we deploy multidimensional mass cytometry combined with unbiased machine learning techniques to immunophenotype peripheral blood mononuclear cells from study participants in cross-sectional and longitudinal cohorts. We show that increases in antigen-experienced adaptive immune cells in the blood, particularly CD45RA-reactivated T effector memory (TEMRA) cells, are associated with early accumulation of brain β-amyloid and with changes in plasma AD biomarkers in still cognitively healthy subjects. Our results suggest that preclinical AD pathology is linked to systemic alterations of the adaptive immune system. These immunophenotype changes may help in the future to identify and develop novel diagnostic tools for early AD assessment and to better understand clinical outcomes.

## INTRODUCTION

Alzheimer’s disease (AD) is a highly prevalent neurodegenerative disorder that leads to progressive memory loss and cognitive as well as functional decline, eventually resulting in dementia symptoms. Accumulation and aggregation of misfolded β-amyloid peptide (Aβ) in the brain represents a primary pathological event of AD **(Jack et al., 2018)** that occurs over decades before symptom onset in a silent, preclinical phase of the disease **(Villemagne et al., 2013)**. Cognitively healthy control subjects (HCS) with elevated brain β-amyloid are individuals at such an early, preclinical stage of the AD continuum, with higher risk for disease progression towards ‘mild cognitive impairment’ (MCI) and established AD dementia stage **(Donohue et al., 2017) (Ossenkoppele et al., 2022)**. Therefore, preclinical AD is currently considered an ideal window of opportunity for disease-modifying therapeutic strategies **(Bateman et al., 2019)**. However, due to a lack of knowledge about prognostic AD biomarkers, it is difficult to identify AD risk populations and populations in which therapeutic interventions might prevent impending neuropathological events. Early disease diagnosis is the subject of intense investigation and in particular, blood-based biomarkers such as isoforms of phosphorylated tau (p-tau181, p-tau217 and p-tau231) as well as Aβ42/Aβ40 ratio in blood plasma have recently been linked to early β-amyloid accumulation in the brain **(Bateman et al., 2020)**. Better characterization of additional systemic changes in the AD continuum would be crucial to improve diagnostic accuracy in the preclinical stage.

The immune system is designed to respond to any homeostatic perturbation, and both innate and adaptive immune cells have been shown to react to AD pathology **(Gaskin et al., 1993) (Monsonego et al., 2003) (Meyer-Luehmann et al., 2008) (Yin et al., 2017) (Rickenbach and Gericke, 2021)**. Moreover, emerging genetic and clinical data indicate an involvement of inflammation-related pathways in AD pathogenesis and disease progression **(Ferretti et al., 2012) (Heneka et al., 2015) (Ransohoff, 2016) (Kunkle et al., 2019)**. Thus, whether immune markers could be used as proxy of β-amyloid accumulation in the brain is a clinically highly relevant question. In this study, we therefore applied state-of-the-art single-cell immunophenotyping, focusing on peripheral T lymphocytes, the largest cellular subtype of the adaptive immune system. After antigen exposure, T cells undergo clonal expansion and differentiation into antigen-specific effector and long-lived memory cells that can spark a faster and more vigorous immune response upon second encounter with their cognate antigen **(Sallusto et al., 1999)**. Priming of the adaptive immune system by brain-derived antigens is still a matter of debate. The existence of protective antibodies against β-amyloid aggregates in healthy aged donors **(Moir et al., 2005) (Sevigny et al., 2016)** strongly indicates that the peripheral immune system can ‘sense’ the brain, most likely via the drainage of brain-derived solutes through dural lymphatic vessels **(Louveau et al., 2015) (Ma et al., 2017) (Rasmussen et al., 2018) (Da Mesquita et al., 2018)**. Human T cells have been shown to be reactive against specific epitopes derived from the linear Aβ1-42 peptide sequence *in vitro* **(Monsonego et al., 2003) (Zota et al., 2009)**. However, while research has focused on the link between T cell reactivity and cognitive outcomes in MCI and AD **(Speciale et al., 2007) (Pellicano et al., 2012) (Gate et al., 2020)**, the association of peripheral T cell alterations with key pathological biomarkers such as cerebral β-amyloid deposition and AD plasma markers in a preclinical stage of AD has not yet been addressed in detail.

Multivariate mass cytometry (CyTOF) combined with data-driven, unbiased analysis tools allows for comprehensive characterization of those complex, disease-associated cellular patterns. The technology has been recently refined **(Brodie and Tosevski, 2017)** and established in cancer biology (e.g. leukemia) **(Levine et al., 2015) (Tsai et al., 2020)**, neurological conditions such as narcolepsy **(Hartmann et al., 2016)** or multiple sclerosis **(Ingelfinger et al., 2022)**, and infectious diseases (e.g. immune factors in *M. tuberculosis* infections) **(Roy Chowdhury et al., 2018)**. To investigate systemic adaptive immune cell alterations during progressing β-amyloidosis across the AD continuum, we employed CyTOF to analyze peripheral blood mononuclear cells (PBMCs) from well characterized cross-sectional and longitudinal subject cohorts. Here, we show that early accumulation of β-amyloid in the brain as well as early changes in plasma AD biomarkers are indeed associated with a systemic increase in effector subpopulations of adaptive immune cells, particularly in T cell lineages, even in the absence of cognitive symptoms. Importantly, we demonstrate that such systemic alterations are independent of typically latent and widespread *Herpesviridae* infections.

## RESULTS

### CyTOF-based Immunophenotyping of Cross-sectional Study Populations Covering the Early AD Spectrum

We analyzed blood-derived material from two separate study populations focusing mainly on the early spectrum of AD, namely cognitively healthy control subjects (‘HCS’) and subjects with mild cognitive impairment (‘MCI’) with biomarker evidence for Alzheimer’s disease. All study participants underwent neuropsychological testing, plasma AD biomarker analysis, and cerebral β-amyloid imaging via positron emission tomography (PET) scanning using [11C]-Pittsburg compound B (PiB) or [18F]-Flutemetamol (FMM) tracers (Figure 1A, Table S1 and S2). To immunophenotype adaptive immune cell populations using CyTOF, we stained live-cell-barcoded PBMCs with self-designed, heavy metal-conjugated antibody panels for the analysis of surface markers on PBMCs, in particular on T cells (Figure 1B). For a first exploratory study (termed ‘*Cross-sectional I*’) we selected PBMCs from 50 individuals (Figure 1C) to represent different cognitive stages of the disease and markedly different β-amyloid status based on PiB PET results. *Cross-sectional I* includes 5 diagnostic groups based on clinical assessment and brain β-amyloid load, with the exception of AD patients, who were derived from a protocol that did not include amyloid PET: ‘AD’ (n = 9), ‘MCI’ (β-amyloid+, n = 9 and β-amyloid–, n = 9), and ‘HCS’ (β-amyloid+, n = 10 and β-amyloid–, n = 13). As expected, cerebral β-amyloid load (PiB SUVR and Centiloid) was significantly different between positive and negative groups (Figure S1A and S1B), with β-amyloid+ groups clearly above the cut-off. Both β-amyloid+ groups were characterized by higher frequency of *APOE* ε4 carriers (approx. 2x in β-amyloid+ compared to β-amyloid– groups) (Figure 1C, Table S1). Furthermore, AD subjects were older than the other groups (p < 0.01, AD versus HCS β-amyloid–) and more likely to be women (Table S1). Since altered levels of plasma AD biomarkers such as Aβ42/Aβ40 ratio **(Schindler et al., 2019)**, p-tau181 **(Janelidze et al., 2020)** and p-tau181/Aβ42 ratio **(Chong et al., 2021)** have been well associated with preclinical AD and MCI due to AD, and are deemed suitable for early diagnosis **(Hansson, 2021)**, we included plasma markers for further characterization (Figure S1C-S1E, Figure 2G). As expected, most significant alterations were observed for HCS β-amyloid+, MCI β-amyloid+ and AD compared to β-amyloid– groups. Plasma total tau, a rough marker of neurodegeneration, was most elevated in AD patients compared to HCS β-amyloid– and MCI β-amyloid+ (Figure S1F).

**Figure 1.**
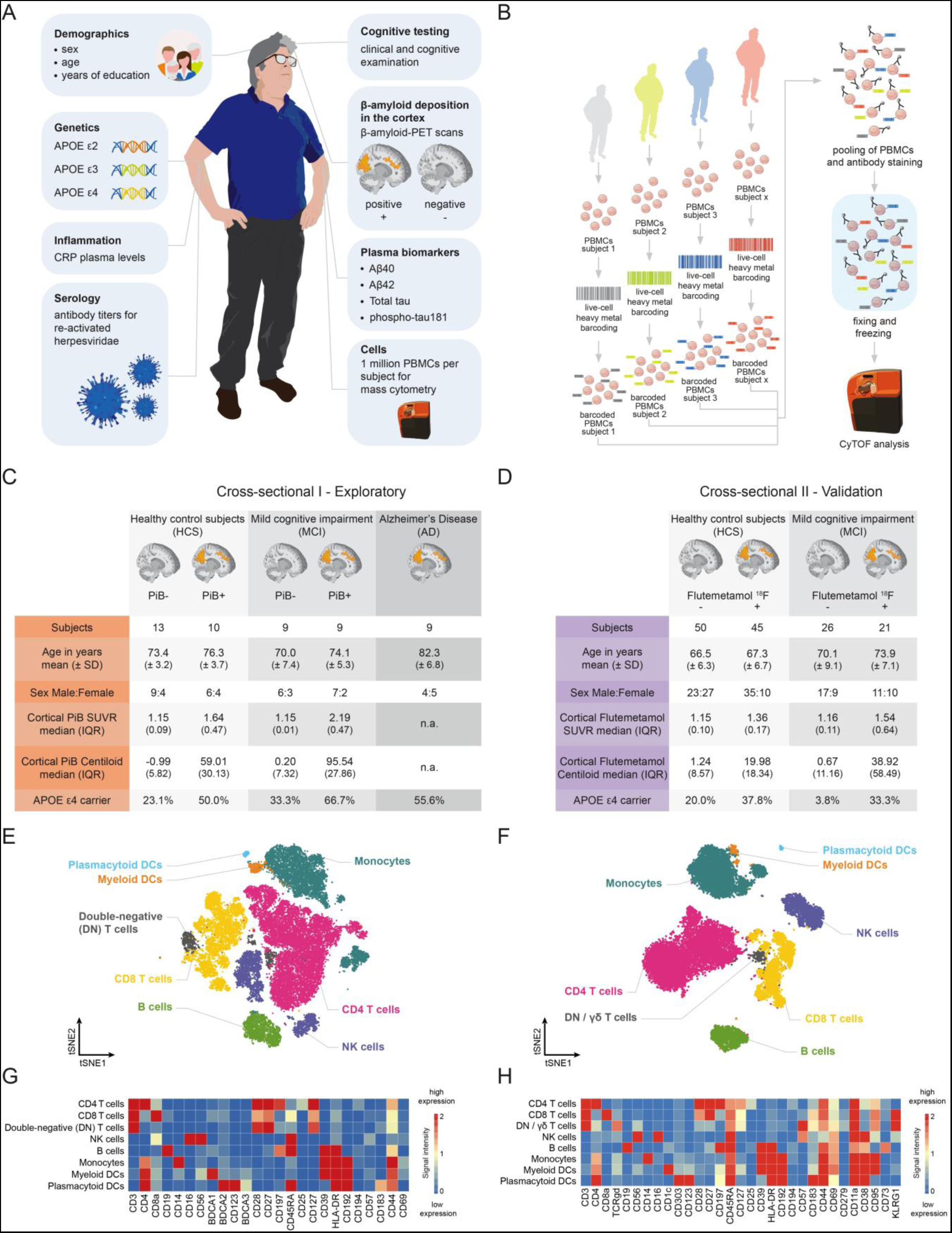
CyTOF-based Immunophenotyping of Cross-sectional Study Populations. **(A)** Overview of available demographic, clinical, and molecular parameters of the subject cohorts in use. **(B)** Schematic workflow: PBMC biobank generated from donor blood, subject-specific barcoding of cells, pooling of barcoded samples into one single bulk sample, application of metal-conjugated antibody staining, fixing and freezing of stained samples, storage till CyTOF acquisition. **(C)** Characteristics of *Cross-sectional I* and its 5 diagnostic groups. Grouping is based on cognitive testing (healthy control subject = HCS, mild cognitive impairment = MCI, Alzheimer’s disease = AD) and cortical β-amyloid measured by [11C]-PiB-PET scans. Subjects are considered β-amyloid positive at a cut-off level of PiB SUVR > 1.265. n.a. = not assessed. **(D)** Characteristics of *Cross-sectional II* and its 4 diagnostic groups. Grouping is based on cognitive testing (healthy control subject = HCS, mild cognitive impairment = MCI) and cortical β-amyloid measured by [18F]-Flutemetamol (FMM)-PET scans. Subjects are considered β-amyloid positive at a cut-off level of FMM Centiloid > 12. **(E, F)** *t*-distributed stochastic neighbor embedding (tSNE) and FlowSOM clustering for all CD45+ PBMCs averaged over all subjects for *Cross-sectional I* **(E)** (tSNE settings: iterations = 12’000, events = 10’000, perplexity = 40; FlowSOM settings: k25, merged into 8 major cell populations) and *Cross-sectional II* **(F)** (tSNE settings: iterations = 4’000, events = 10’000, perplexity = 50; FlowSOM settings: k15, merged into 8 major cell populations). **(G, H)** Heatmap for identification of 8 major CD45^+^ PBMC clusters. Heatmap shows *arcsinh*-transformed median marker intensities for *Cross-sectional I* **(G)** and *Cross-sectional II* **(H)**.

**Figure 2.**
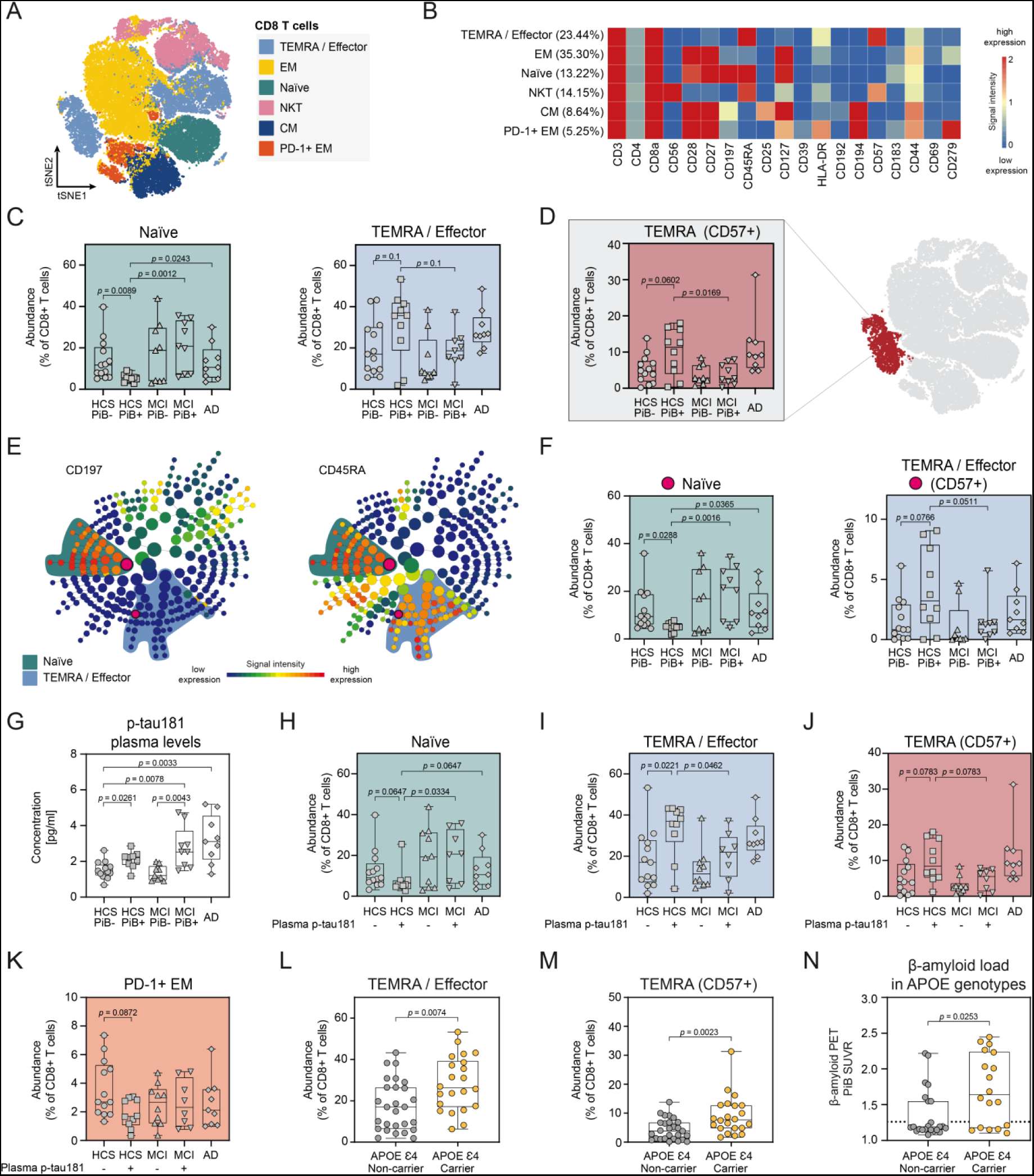
*Cross-sectional I* - Increase of Peripheral CD8^+^ TEMRA/Effector Cells in AD-biomarker-positive Cognitively Healthy Individuals. **(A)** tSNE and FlowSOM clustering for PBMC-derived CD8^+^ T cells averaged over all subjects of *Cross-sectional I* (tSNE settings: iterations = 2’000, events = max. 10’000, perplexity = 50; FlowSOM settings: k17, merged into 6 major clusters). EM = effector memory cells, NKT = natural killer T cells, CM = central memory cells. **(B)** Heatmap for identification of 6 major CD8^+^ T cell clusters. Heatmap shows *arcsinh*-transformed median marker intensities. **(C)** Quantification and comparison of relative abundances of CD8^+^ T cell subpopulations in HCS PiB+ subjects with HCS PiB–, MCI PiB+ and AD patients. Naïve, non-activated CD8^+^ T cells are decreased in HCS PiB+ individuals compared to HCS PiB–, MCI PiB+ and AD groups. CD8^+^ TEMRA/Effector cells in HCS PiB+ tend to be increased compared to HCS PiB– and MCI PiB+ subjects. **(D)** Analysis of subclusters of CD8 TEMRA/Effector cells indicates significant changes in CD57^+^ terminally differentiated TEMRA cells. CD57^+^ TEMRA cells are elevated in HCS PiB+ individuals compared to HCS PiB– and MCI PiB+ subjects. **(E)** CITRUS-generated maps showing CD8^+^ T cell clusters with differential abundance (marked with pink dot). Cell cluster size relates to the dimension of the depicted sphere. Representative graphs show CD197 (CCR7) and CD45RA median marker intensity per cluster. Significantly different subclusters are located in naïve (CCR7^+^/CD45RA^+^) and TEMRA/Effector (CCR7^-^/CD45RA^+^) CD8^+^ T cell clusters. **(F)** CITRUS-based cluster quantification for specific subpopulations of CD8^+^ T cells. SAM analysis reveals reduced relative abundance of naïve CD8^+^ T cells in HCS PiB+ compared to HCS PiB–, MCI PiB+ and AD patients. Relative abundance of a CD8^+^ CD57^+^ subpopulation of TEMRA/Effector T cells is increased in HCS PiB+ individuals compared to HCS PiB– and MCI PiB+. **(G)** Plasma AD biomarker quantification shows increased levels of p-tau181 in HCS PiB+ vs. HCS PiB–, MCI PiB+ vs. MCI PiB–, MCI PiB+ vs. HCS PiB– and AD vs. HCS PiB–. **(H-J)** HCS and MCI groups were divided into plasma p-tau181-low (–) and p-tau181-high (+) subgroups based on p-tau181 median split at 1.89 pg/ml. Relative abundance of naïve CD8^+^ T cells **(H)** is decreased in HCS+ individuals compared to HCS–, MCI+ and AD groups. CD8^+^ TEMRA/Effector cells **(I)** and CD57^+^ TEMRA cells **(J)** are elevated in HCS+ compared to HCS– and MCI+ subjects. **(K)** Cluster containing CD8^+^ PD-1^+^ EM with lower relative abundance in HCS+ (p-tau181-high) compared to HCS–. **(L-N)** Relative abundances of CD8^+^ TEMRA/Effector cells **(L)** and CD57^+^ TEMRA cells **(M)** are elevated in *APOE* ε4 carriers compared to non-carriers. Potential confounding effect explained by significantly higher median β-amyloid load in *APOE* ε4 carriers **(N)**. Dotted line represents PiB SUVR cut-off = 1.265. Each symbol in quantification graphs represents data from one subject. Box plot represents interquartile range (IQR), horizontal line indicates median, whiskers extend to min/max data point. The *p* values were calculated via nonparametric Mann-Whitney (L-N) or Kruskal-Wallis tests (C, D, F-K). Controlling for multiple comparisons was performed with false-discovery-rate (FDR) method of Benjamini and Hochberg. Selected *p* values ≤ 0.1 (FDR 10%) are displayed.

A second, independent cross-sectional study (termed ‘*Cross-sectional II’*) was analyzed to extend and reproduce key findings of *Cross-sectional I*. Here, the study population consisted of 142 participants including 4 diagnostic groups: HCS (β-amyloid+, n = 45 and β-amyloid–, n = 50) and MCI (β-amyloid+, n = 21 and β-amyloid–, n = 26) (Figure 1D). In this study, subjects with FMM SUVR and Centiloid more closely around the cut-off were included, resulting in a more homogeneous distribution of cerebral β-amyloid load among these diagnostic groups compared to *Cross-sectional I*. Within the β-amyloid+ MCI groups, this is reflected by a much lower median FMM Centiloid = 38.92 (IQR = 58.49) in *Cross-sectional II* compared to PiB Centiloid = 95.54 (IQR = 27.86) in *Cross-sectional I* (Figure S1K). Still, cerebral β-amyloid load (FMM SUVR and Centiloid) was significantly different between β-amyloid+ and β-amyloid– groups (Figure S1H and S1I). Both β-amyloid+ groups had a higher frequency of *APOE* ε4 carriers (Figure 1D, Table S2).

After CyTOF acquisition, obtained datasets were depicted via automated dimensionality reduction tools (here: tSNE) and analyzed with data-driven, unbiased clustering algorithms (here: FlowSOM) (Figure 1E and 1F). The resulting clusters were compared to heatmaps, which contain the median signal intensities for every marker of interest (Figure 1G and 1H) and allow for assignment of clusters to biologically meaningful cell types. Automated clustering successfully identified major CD45^+^ PBMC immune cell populations such as CD4^+^ and CD8^+^ T cells, double-negative (DN) / γδ T cells, CD19^+^ B cells, CD14^+/low^ CD16^+/–^ monocytes, CD16^+/low^ CD56^+^ natural killer (NK) cells and blood-derived dendritic cells (DCs), which split into BDCA1^+^ myeloid DCs (mDCs) and BDCA2^+^ plasmacytoid DCs (pDCs) (Figure 1E-1H). Adaptive immune cell populations were largely comparable among the different diagnostic groups (Figure S1G and S1J). Of note, the relative abundance of total CD8^+^ T cells was slightly increased in HCS β-amyloid+ subjects compared to AD samples in *Cross-sectional I* (Figure S1G). In the following, we took further insight into adaptive immune cell populations by subclustering T cell subpopulations of interest for in-depth analysis.

### Cross-sectional I *-* Increase of Peripheral CD8^+^ TEMRA/Effector Cells in AD-biomarker-positive Cognitively Healthy Individuals

CD3^+^ CD4^−^ CD8^+^ T cells were preselected and analyzed via FlowSOM automated clustering (Figure 2A). Six major CD8^+^ T cell subpopulations were identified via median signal intensities of lineage and activation markers (Figure 2B). One of the most prominent CD8^+^ T cell subclusters was a mixed cluster of CD197^−^ CD45RA-reactivated T effector memory cells (‘TEMRA’ cells) and T effector cells (together termed ‘TEMRA/Effector’ cells) (Median = 23.4% of CD8^+^ T cells; IQR = 21.2%) **(Sallusto et al., 2004)**. Moreover, we found CD197^+^ CD45RA^+^ naïve T cells, CD56^+^ NK T cells, CD197^+^ CD45RA^−^ central memory (CM) T cells, CD197^−^ CD45RA^−^ effector memory (EM) T cells, as well as a potentially exhausted variant of EM T cells (HLA-DR^+^ PD-1^+^).

Relative abundances of CD8^+^ T cell subpopulations were compared between an HCS β-amyloid+ group of interest (= preclinical AD, positive for β-amyloid PiB PET) and HCS β-amyloid–, MCI β-amyloid+ as well as AD patients (Figure 2C). Statistical analysis indicated that relative abundance of naïve CD8^+^ T cells was significantly reduced in HCS β-amyloid+ study participants compared to HCS β-amyloid–, MCI β-amyloid+ and AD subjects. In contrast, CD8^+^ TEMRA/Effector cells showed only a tendency to increase in HCS β-amyloid+ study participants compared to HCS β-amyloid– and MCI β-amyloid+ subjects (*p* = 0.1). The CD8^+^ TEMRA/Effector cell population is a heterogeneous group; therefore, in order to characterize its increase in HCS β-amyloid+ individuals in greater detail, we subclustered the TEMRA/Effector cell cluster and compared cell abundances across groups. We found that a terminally differentiated, reactivated and highly antigen-specific TEMRA subpopulation, CD57^+^ TEMRA cells **(Brenchley et al., 2003) (Mahnke et al., 2013) (Verma et al., 2017)**, was increased in HCS β-amyloid+ subjects compared to HCS β-amyloid– and MCI β-amyloid+ (Figure 2D). The sex of the study participants did not affect immune cell clusters of interest (Pearson’s chi-squared test: *p*(CD8^+^ naïve T cells) = 1.00; *p*(CD8^+^ TEMRA/Effector cells) = 1.00; *p*(CD8^+^ CD57^+^ TEMRA cells) = 1.00), despite the imbalanced male/female distribution in *Cross-sectional I* (Figure 1C, Table S1). Age differences among diagnostic groups had only minor effect on CD8^+^ T cells’ relative cluster abundances (Figure S2A, Spearman correlation: -0.5 < rho < 0.5). Additional multiple linear regression analysis for age and sex was consistent with these results, with the only difference being that naïve CD8^+^ T cells were significantly affected by age, but only with low estimate close to 0 (Figure S2C).

We confirmed our findings using an alternative clustering and analysis via CITRUS and its association models, which enables a fully automated discovery of statistically significant biological signatures within single-cell datasets **(Bruggner et al., 2014)**. The most significantly different clusters resulting from CITRUS analysis in combination with the correlative SAM association model were highlighted in hierarchical clustering maps (Figure 2E). Marker expression analysis revealed that these cluster subsets belonged to naïve CD8^+^ T cells as well as a subpopulation of CD57^+^ TEMRA/Effector cells. Quantification of these clusters revealed that results on naïve and TEMRA/Effector CD8^+^ T cells could be replicated using correlative CITRUS analysis (Figure 2F).

In an alternative diagnostic grouping, HCS and MCI subjects were subdivided according to median split of plasma p-tau181 levels (1.89 pg/ml) instead of β-amyloid PiB PET. For readability, we refer to subjects who are above the median as ’p-tau181+’, being aware that this is not a clinical cut-off for tau positivity. Relative abundance of CD8^+^ naïve T cells was reduced in HCS p-tau181+ subjects compared to HCS p-tau181–, MCI p-tau181+ and AD, whereas TEMRA/Effector and CD57^+^ TEMRA cells were increased in HCS p-tau181+ study participants in comparison with HCS p-tau181– and MCI p-tau181+ (Figure 2G-2J). Thus, using two independent clustering methods as well as two different AD biomarkers, we show that a preclinical AD stage with early cerebral β-amyloidosis in cognitively healthy subjects (HCS β-amyloid+/p-tau181+) is characterized by an increase in CD8^+^ TEMRA/Effector cells with a concomitant decrease in CD8^+^ naïve T cells. In addition, specifically when we analyzed plasma p-tau181, we found a decrease in PD-1^+^ (CD279^+^), potentially exhausted EM cells in HCS p-tau181+ compared to HCS p-tau181– (Figure 2K).

To control for association with *APOE* ε4 genotype, the most prominent risk factor for AD, we compared relative abundances of our immune populations of interest among *APOE* ε4 carriers and non-carriers. We found that CD8^+^ TEMRA/Effector and CD57^+^ TEMRA cells were elevated in *APOE* ε4 carriers compared to non-carriers (Figure 2L and 2M). However, *APOE* ε4 genotype might be a confounding variable in this case because we observed, as expected, significantly higher median β-amyloid load in *APOE* ε4 carriers (Figure 2N).

### β-Amyloid-positive, Cognitively Healthy Individuals Are Further Characterized by Reduced Naïve CD4^+^ T Cell Numbers

CD3^+^ CD4^+^ CD8^−^ cells were preselected, clustered and identified as shown before (Figure S3A and S3B). Similar to what we observed for CD8^+^ T cells, relative abundance of CD4^+^ naïve T cells was lower in HCS β-amyloid+ individuals compared to MCI β-amyloid+ and AD patients (Figure S3C). However, the majority of other CD4^+^ T cell subclusters did not exhibit statistically significant differences in relative abundances. Only CD4^+^ effector memory T cells were increased in HCS β-amyloid+ subjects compared to AD patients. Age differences between diagnostic groups had no effect on relative cluster abundances of CD4^+^ T cells (Figure S2B, Spearman correlation: -0.5 < rho < 0.5). The sex of the study participants did not affect CD4^+^ T cell clusters of interest (Pearson’s chi-squared test: *p*(CD4^+^ naïve T cells) = 1.00; *p*(CD4^+^ EM cells) = 1.00). No differences in CD4^+^ T cell clusters were observed when diagnostic groups were formed based on plasma AD biomarkers.

### Cross-sectional II - Confirmation of Increased CD8^+^ TEMRA/Effector Cell Abundance in Early Stages of AD

To specifically investigate the reproducibility and robustness of the prominent CD8^+^ T cell results from *Cross-sectional I*, we analyzed a larger study population, termed ‘*Cross-sectional II*’. The selection of the participant was less stringent on β-amyloid status and also included values closer to the cut-off compared to *Cross-sectional I*. Due to updated marker panels, we were able to conduct a more detailed cluster characterization and identification via tSNE and FlowSOM clustering, identifying 7 major subpopulations (Figure 3A and 3B). We compared the median relative abundance of CD8^+^ T cell clusters across diagnostic groups. Based on a false discovery rate of 10%, we found a significant decrease in CD8^+^ naïve T cells and an increase in CD8^+^ TEMRA/Effector cells in HCS β-amyloid+ and, interestingly, also in MCI β-amyloid+ subjects compared to HCS β-amyloid– (Figure 3C). Age differences between diagnostic groups had only minor effect on relative cluster abundances (Figure S2D, Spearman correlation: -0.5 < rho < 0.5). Although the male/female distribution in *Cross-sectional II* was imbalanced (Figure 1D, Table S2), the sex of the study participants had no influence on immune cell clusters of interest (Pearson’s chi-squared test: *p*(CD8^+^ naïve T cells) = 0.50; *p*(CD8^+^ TEMRA/Effector cells) = 1.00). Additional multiple linear regression analysis confirmed again that CD8^+^ T cell cluster abundances were largely independent of age and sex; only naïve CD8^+^ T cells were significantly affected by age, albeit with low estimate close to 0 (Figure S2E). We found that CD8^+^ TEMRA/Effector cells were elevated in *APOE* ε4 carriers compared to non-carriers (Figure 3D). Also here, *APOE* ε4 genotype is a confounding variable since we observed significantly higher median β-amyloid load in *APOE* ε4 carriers (Figure 3E).

**Figure 3.**
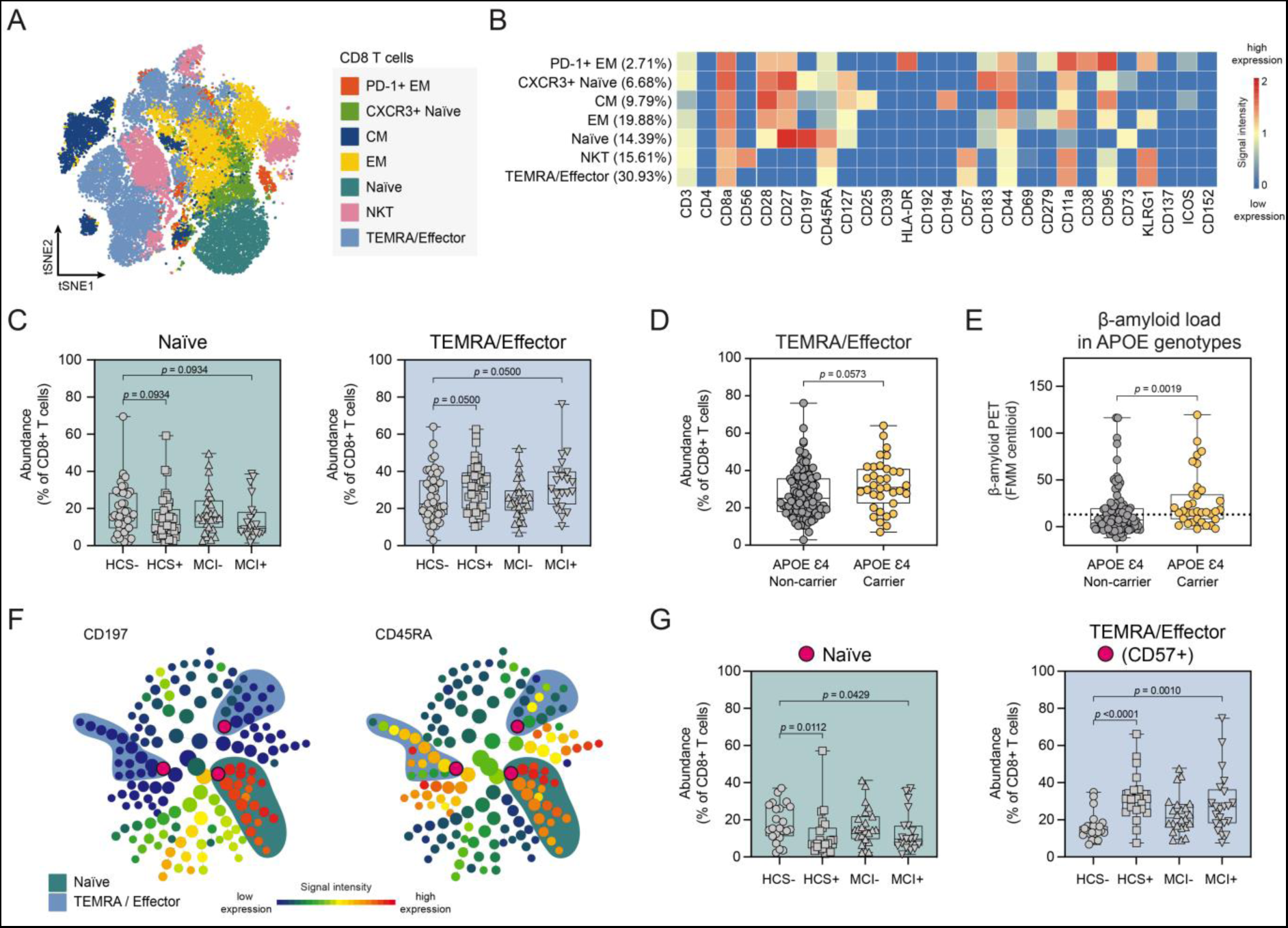
*Cross-sectional II* - Confirmation of Increased CD8^+^ TEMRA/Effector Cell Abundance in Early Stages of AD. **(A)** tSNE and FlowSOM clustering for PBMC-derived CD8^+^ T cells averaged over all subjects of *Cross-sectional II* (tSNE settings: iterations = 4’000, events = max. 10’000, perplexity = 50; FlowSOM settings: k30, merged into 7 major clusters). EM = effector memory cells, NKT = natural killer T cells, CM = central memory cells. **(B)** Heatmap for identification of 7 major CD8^+^ T cell clusters. Heatmap shows *arcsinh*-transformed median marker intensities. **(C)** Quantification and comparison of relative abundances of CD8^+^ T cell subpopulations in HCS β-amyloid– subjects with HCS β-amyloid+ and MCI β-amyloid+ subjects. Naïve CD8^+^ T cells are reduced in both HCS β-amyloid+ and MCI β-amyloid+ individuals compared to HCS β-amyloid–. CD8^+^ TEMRA/Effector cells in HCS β-amyloid+ and MCI β-amyloid+ show higher abundance compared to HCS β-amyloid– subjects. **(D, E)** Relative abundance of CD8^+^ TEMRA/Effector cells is elevated in *APOE* ε4 carriers compared to non-carriers **(D)**. Potential confounding effect explained by significantly higher median β-amyloid load in *APOE* ε4 carriers **(E)**. Dotted line represents FMM Centiloid cut-off = 12. **(F)** Cluster maps generated with CITRUS for CD8+ T cells from 90 randomly selected samples show cell clusters with differential abundance (marked with pink dot). Representative graphs show CD197 (CCR7) and CD45RA median marker intensity per cluster. Significantly changed clusters reside in naïve (CCR7^+^/CD45RA^+^) and TEMRA/Effector (CCR7^-^/CD45RA^+^) CD8^+^ T cell clusters. **(G)** CITRUS-based cluster quantification (correlative SAM model) reveals reduced relative abundance of naïve CD8^+^ T cells in HCS β-amyloid+ and MCI β-amyloid+ individuals compared to HCS β-amyloid–. CD8^+^ CD57^+^ TEMRA/Effector T cells are increased in HCS β-amyloid+ and MCI β-amyloid+ subjects compared to HCS β-amyloid–. Each symbol in quantification graphs represents data from one subject. Box plot represents IQR, horizontal line indicates median, whiskers extend to min/max data point. The *p* values were calculated via nonparametric Mann-Whitney (D, E) or Kruskal-Wallis tests (C, G). Controlling for multiple comparisons was performed with FDR method of Benjamini and Hochberg. Selected *p* values ≤ 0.1 (FDR 10%) are displayed.

We confirmed our findings using correlative CITRUS clustering. Hierarchical cluster maps generated with CITRUS for CD8^+^ T cells from 90 randomly selected samples show cell clusters with differential abundance (Figure 3F). Both CD8^+^ naïve T cells and CD8^+^ CD57^+^ TEMRA/Effector cells were significantly altered in HCS β-amyloid+ and MCI β-amyloid+ subjects compared to HCS β-amyloid– (Figure 3G).

Thus, we confirmed in a second independent analysis that a preclinical AD stage with early cerebral β-amyloidosis (HCS β-amyloid+) is accompanied by alterations in the peripheral T cell compartment. Importantly, in *Cross-sectional II*, also in the ‘MCI due to AD’ stage (MCI β-amyloid+) the above-mentioned immune alterations can be observed. Since median β-amyloid load in the MCI β-amyloid+ group of *Cross-sectional II* was lower than in the HCS β-amyloid+ group of *Cross-sectional I* (Figure S1K), the extent of β-amyloid-pathology is less advanced in these subjects and more comparable to the HCS β-amyloid+ stage of *Cross-sectional I*. Therefore, in *Cross-sectional II*, T cell alterations are observable in both β-amyloid+ healthy and MCI subjects.

### β-Amyloid-associated Effects on Peripheral Adaptive Immune Cells Are Independent of Latent Herpesvirus Infections

The results of both cross-sectional studies indicate that early β-amyloid accumulation in the brain is characterized by an altered immune profile in the blood, suggesting a link between brain β-amyloidosis and the peripheral immune system. However, certain diagnostic groups could have been biased by subclinical levels of viral infections. In particular, herpesvirus infections have been suggested to be associated with AD pathogenesis **(Ball, 1982)**, and might be a driver of the observed peripheral immune alterations. To study this, we tested for possible associations between our predefined diagnostic groups and serum antibody titers against *Herpesviridae* such as VZV, HSV-1, HSV-2, EBV and CMV. We examined IgM as well as IgG titers, to determine both acute infection and long-term immunity, respectively (Figure S4A and S4B). All study participants proved to be above the diagnostic threshold for various *Herpesviridae* IgG. Of note, 46% of subjects in *Cross-sectional I* and 40% of subjects in *Cross-sectional II* were tested positive for anti-CMV IgG, a virus known to affect the proportions of peripheral CD8^+^ TEMRA cells **(Weekes et al., 1999)**. In contrast, only very few cases with positive IgM titers were observed in both cohorts, indicating low levels of acute infections. Statistical testing revealed no significant differences in antiviral IgG serum titers between diagnostic groups, indicating that *Herpesviridae* infections did not preferentially occur in a specific cognitive or β-amyloid state (Figure 4A and 4F).

**Figure 4.**
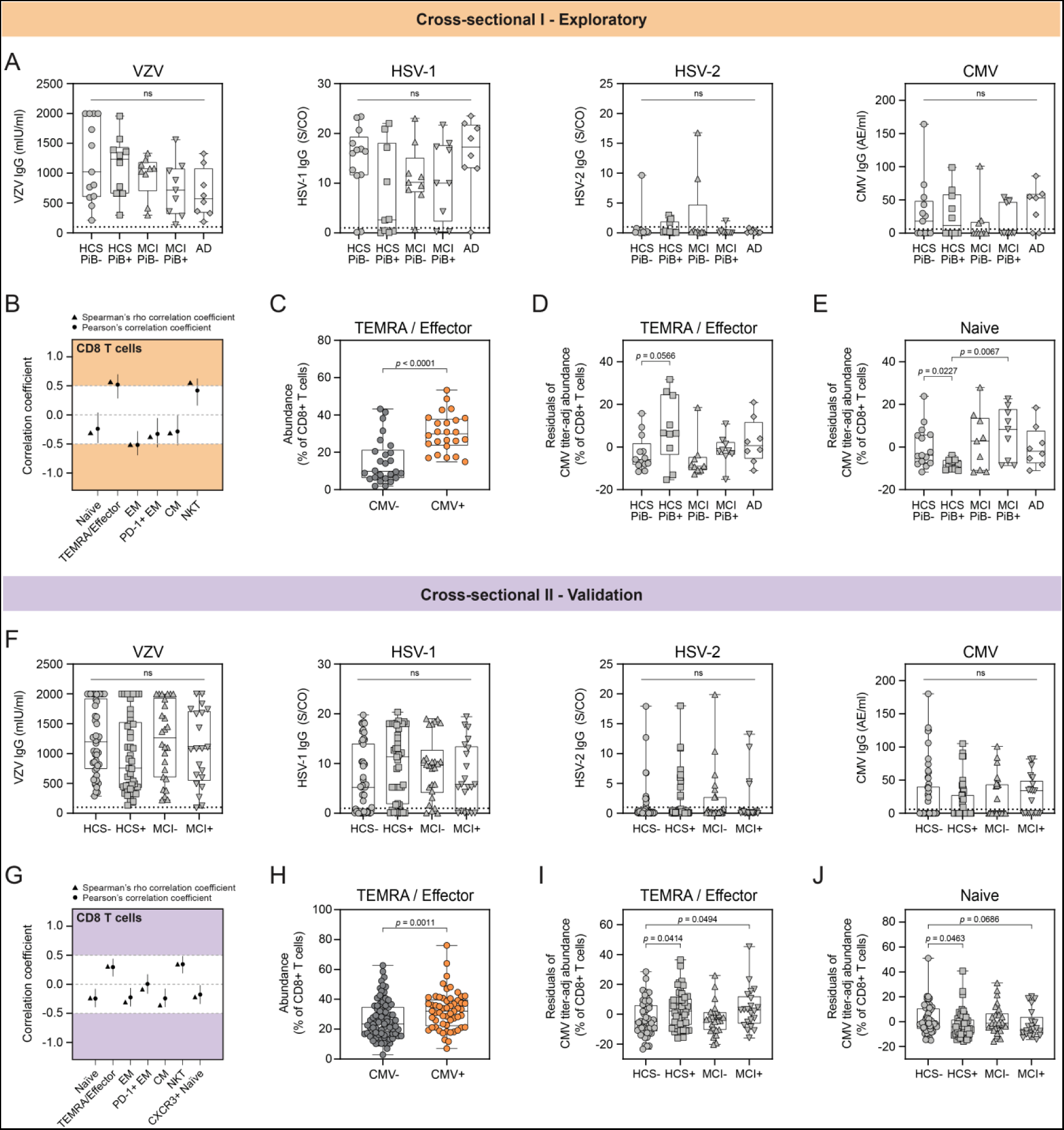
β-Amyloid-associated effects on peripheral adaptive immune cells are independent of latent herpesvirus infections. *Cross-sectional I*: **(A)** Quantification of IgG titers against VZV (in mIU/ml, threshold >100), HSV-1 (in S/CO, threshold >1), HSV-2 (in S/CO, threshold >1), and CMV (in AE/ml, threshold ≥6) for every subject in each of the diagnostic groups. Displayed are the absolute IgG antibody titers. Dotted lines mark reference diagnostic thresholds. Statistical testing shows no significant (ns) difference between diagnostic groups. **(B)** Correlations between CMV IgG titers of all subjects and relative CD8^+^ T cell cluster abundance. Black dots represent the estimated Pearson’s r correlation coefficients and the black lines the corresponding 95% confidence intervals. Black triangles represent the estimated Spearman’s rho correlation coefficients. CD8^+^ T cell subclusters such as TEMRA/Effector cells, EM and NKT cells appear to be affected by changing CMV IgG titers. **(C)** Individuals which have been tested positive for CMV IgG (CMV+) exhibit a significantly higher relative abundance of CD8^+^ TEMRA/Effector cells compared to CMV– subjects. **(D, E)** Statistical testing for adjusted residuals reveals that CMV IgG titer-adjusted CD8^+^ TEMRA/Effector cell abundances are significantly elevated in HCS PiB+ individuals compared to HCS PiB– **(D)**. CD8^+^ naïve cell abundance is reduced in HCS PiB+ subjects compared to HCS PiB– and MCI PiB+ **(E)**. *Cross-sectional II*: **(F)** Quantification of IgG titers against VZV, HSV-1, HSV-2, and CMV for every subject in each of the diagnostic groups. Displayed are the absolute IgG antibody titers. Dotted lines mark reference diagnostic thresholds. Statistical testing shows no significant (ns) difference between diagnostic groups. **(G)** Correlations between CMV IgG titers and relative CD8^+^ T cell cluster abundance. **(H)** CMV+ subjects show significantly higher relative abundance of CD8^+^ TEMRA/Effector cells compared to CMV–. **(I, J)** Statistical testing for adjusted residuals reveals that CMV IgG titer-adjusted CD8^+^ TEMRA/Effector cell abundances are significantly elevated in HCS β-amyloid+ and MCI β-amyloid+ individuals compared to HCS β-amyloid– **(I)**. CD8^+^ naïve cell abundance is reduced in HCS β-amyloid+ and MCI β-amyloid+ subjects compared to HCS β-amyloid– **(J)**. Each symbol in quantification graphs represents data from one subject. Box plot represents IQR, horizontal line indicates median, whiskers extend to min/max data point. The *p* values were calculated using nonparametric Mann-Whitney (C, H) or Kruskal-Wallis tests (A, D, E, F, I, J). Controlling for multiple comparisons was performed with FDR method of Benjamini and Hochberg. Selected *p* values ≤ 0.1 (FDR 10%) are displayed.

We next checked for specific influences of *Herpesviridae* on CD8^+^ T cell subpopulations by correlating relative cluster abundances of FlowSOM-derived subclusters with serum antibody titers of the four quantitatively assessed *Herpesviridae* (VZV, CMV, HSV-1, HSV-2). In both cross-sectional studies, none of the CD8^+^ T cell subclusters were substantially affected by increasing IgG titers against VZV, HSV-1 and HSV-2 (Figure S4C and S4D, Spearman correlation: -0.5 < rho < 0.5). However, CD8^+^ T cell subclusters were affected by altered anti-CMV IgG titers mainly in *Cross-sectional I*, less so in *Cross-sectional II* (Figure 4B and 4G). Specifically, CD8^+^ EM T cells were negatively correlated with increasing anti-CMV IgG levels (rho ≤ -0.5), whereas CD8^+^ TEMRA/Effector and NK T cells were positively correlated (rho ≥ 0.5) (Figure 4B). No differences in anti-CMV IgG titers have been found in cerebral β-amyloid groups and APOE genotypes (Figure S4E and S4F).

As expected, we observed that individuals who had been tested positive for anti-CMV IgG (CMV+) exhibited a significantly higher relative abundance of CD8^+^ TEMRA/Effector cells compared to CMV– subjects (Figure 4C and 4H). As this was a cell population we identified as characteristic signature cluster associated with preclinical β-amyloidosis, we adjusted the relative abundances of our CD8^+^ T cell clusters of interest for anti-CMV IgG titer levels using linear regression to control for the effect of CMV infection. Statistical testing for the adjusted residuals revealed that anti-CMV IgG titer-adjusted CD8^+^ TEMRA/Effector cells were still significantly elevated in HCS β-amyloid+ individuals compared to HCS β-amyloid– subjects in both *Cross-sectional I* and *II* (Figure 4D and 4I). Additionally, in *Cross-sectional II*, CD8^+^ TEMRA/Effector cells were still increased in MCI β-amyloid+ patients compared to HCS β-amyloid– (Figure 4I). Significant decreases in naïve CD8^+^ T cells also remained largely unchanged after CMV titer adjustment (Figure 4E and 4J).

Thus, we confirmed that the observed immune cell signature clusters of CD8^+^ T cells, which characterize a preclinical stage of early β-amyloidosis in cross-sectional setups, are independent of latent, reactivated or acute *Herpesviridae* background infections. No significant association was found between herpesvirus IgG titers and CD4^+^ T cell clusters from *Cross-sectional I*.

### Peripheral Immune Profiling for AD Biomarker-accumulating Subjects: A Longitudinal Study Design

After identifying an immune pattern associated with early β-amyloid deposition in the brain of cognitively healthy subjects, we asked whether these changes were also related to AD biomarker accumulation over time. For that reason, we investigated a cohort of 59 elderly, cognitively unimpaired subjects who were followed over a time period of three years. For each individual we assessed the cerebral β-amyloid load via two [11C]-PiB-PET scans at baseline and after 3 years at the end of the study and collected blood for PBMC isolation at *Baseline*, after 1.5 years (termed ‘*Follow-up1*’) and after 3 years (Termed ‘*Follow-up2*’) (Figure 5A). Plasma AD biomarkers such as p-tau181 were assessed at *Baseline* and *Follow-up2*.

**Figure 5.**
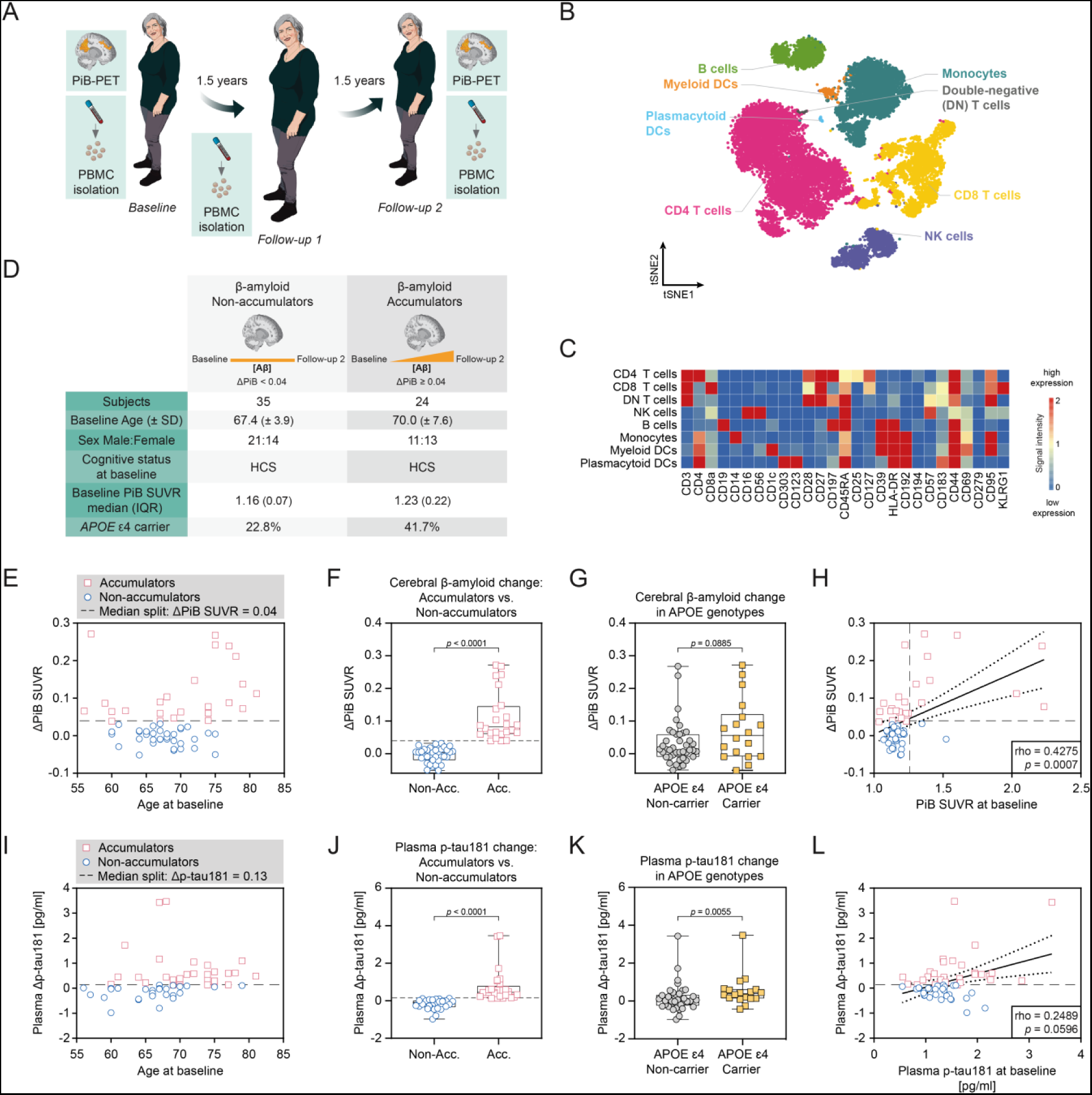
Peripheral Immune Profiling for AD Biomarker-accumulating Subjects: A Longitudinal Study Design. **(A)** Longitudinal study participants were cognitively healthy at baseline and donated blood three times over a period of 3 years: (i) at *Baseline*, (ii) at *Follow-up1* and (iii) at *Follow-up2*. All individuals received two [11C]-PiB-PET scans at *Baseline* and *Follow-up2*. **(B)** tSNE and FlowSOM clustering for all CD45^+^ PBMCs of all subjects at all available timepoints (tSNE settings: iterations = 2’000, events = 10’000, perplexity = 40; FlowSOM settings: k25, merged into 8 major cell populations). **(C)** Heatmap for identification of 8 major CD45^+^ PBMC clusters. Heatmap shows *arcsinh*-transformed median marker intensities. **(D-G)** Group allocation into cortical β-amyloid Non-accumulators and Accumulators over time. Overview of demographic, clinical and genetic characteristics in respective groups **(D)**. Median split at ΔPiB SUVR = 0.04 is used to define β-amyloid Non-accumulators (ΔPiB SUVR < 0.04, blue circles) and β-amyloid Accumulators (ΔPiB SUVR ≥ 0.04, red rectangles) **(E)**. ΔPiB SUVR is significantly higher in β-amyloid Accumulators **(F)** and *APOE* ε4 carriers **(G)**. ΔPiB SUVR = PiB SUVR(*Follow-up2*) – PiB SUVR(*Baseline*). Horizontal dashed line represents cut-off for β-amyloid Accumulators. **(H)** Correlation of ΔPiB SUVR with PiB SUVR at baseline with linear regression; horizontal dashed line represents cut-off for β-amyloid Accumulators (ΔPiB SUVR ≥ 0.04); vertical dashed line represents cut-off for cerebral β-amyloid-positive subjects (PiB SUVR = 1.223). **(I-K)** Alternative group allocation based on plasma p-tau181 changes over time. Median split at Δp-tau181 = 0.13pg/ml is used to define p-tau181 Non-accumulators (Δp-tau181 < 0.13, blue circles) and p-tau181 Accumulators (Δp-tau181 ≥ 0.13, red rectangles) **(I)**. Δp-tau181 is significantly higher in Accumulators **(J)** and *APOE* ε4 carriers **(K)**. Δp-tau181 = p-tau181(*Follow-up2*) – p-tau181(*Baseline*). Horizontal dashed line represents cut-off for p-tau181 Accumulators. **(H)** Correlation of Δp-tau181 with p-tau181 levels at baseline with linear regression; horizontal dashed line represents cut-off for p-tau181 Accumulators (Δp-tau181 ≥ 0.13). Each symbol in quantification graphs represents data from one subject. Box plot represents IQR, horizontal line indicates median, whiskers extend to min/max data point. The *p* values were calculated via nonparametric Mann-Whitney tests (5F, 5G and 5J, 5K). All *p* values ≤ 0.1 (FDR 10%) are displayed. Correlation coefficients (5H, 5L) were calculated via Spearman approach (rho and *p* value reported). Dotted lines represent 95% confidence intervals of a linear regression.

We analyzed PBMCs of all available time points (Table S3) from all 59 study participants with improved versions of CyTOF antibody panels for the characterization of T cells. After CyTOF acquisition, datasets were clustered and visualized in tSNE/FlowSOM maps (Figure 5B). Single marker signal intensities per cluster were used to identify the major clusters of human PBMCs (Figure 5C). Since we were interested in understanding the association of AD pathology, particularly cerebral β-amyloid accumulation and plasma p-tau181 increase, with peripheral immune cell signatures, we divided subjects into groups of individuals whose cerebral β-amyloid or p-tau181 levels remained largely stable over time (termed ‘Non-accumulators’) and those whose levels increased over time (termed ‘Accumulators’). For β-amyloid analysis, based on the change in cortical PiB SUVR over time (ΔPiB SUVR = PiB SUVR*Follow-up2* – PiB SUVR*Baseline*), we split all ΔPiB SUVR values ≥ 0 by median to categorize individuals with ΔPiB SUVR ≥ 0.04 as β-amyloid Accumulators (n = 24) and individuals with ΔPiB SUVR < 0.04 as β-amyloid Non-accumulators (n = 35) (Figure 5D and 5E). Median ΔPiB SUVR for Accumulators (ΔPiB SUVR = 0.082, IQR = 0.078) was significantly higher compared to Non-accumulators (ΔPiB SUVR = 0.003, IQR = 0.028) (Figure 5F). Therefore, this approach allowed us to identify only Accumulators with clearly increasing PiB retention over time compared to Non-accumulators. Correlation of ΔPiB SUVR with each individual’s age at *Baseline* revealed no strong association over time (Figure S5A, rho = 0.2287, *p* = 0.0814). As expected, *APOE* ε4 carriers had a higher median ΔPiB SUVR compared to non-carriers (Figure 5G and S5B). Out of 35 Non-accumulators, 2 subjects were β-amyloid+ at *Baseline*, whereas out of 24 Accumulators, 14 subjects were β-amyloid+ at *Baseline* (Figure S5B). Moreover, we observed a significant correlation between high *Baseline* PiB SUVR and increased ΔPiB SUVR over time (Figure 5H, rho = 0.4275, *p* = 0.0007).

Alternatively, subjects were categorized according to the change of plasma p-tau181 levels over time (Δp-tau181). Plasma p-tau181 levels were split by median at Δp-tau181 = 0.13pg/ml, to distinguish between p-tau181 Accumulators and Non-accumulators (Figure 5I). Plasma Δp-tau181 was significantly higher in p-tau181 Accumulators and *APOE* ε4 carriers (Figure 5J and 5K). Higher *Baseline* p-tau181 levels were not overly associated with increasing Δp-tau181 (Figure 5L, rho = 0.2489, *p* = 0.0596). In contrast, age at *Baseline* was significantly correlated with higher plasma Δp-tau181 (Figure S5C, rho = 0.4881, *p* = 0.0001).

### Longitudinal Analysis Indicates CD8^+^ T Cell Alterations in AD Biomarker Accumulators

CD3^+^ CD4^-^ CD8^+^ T cells were subclustered via FlowSOM (Figure S6A). Without multiple comparison analysis, we increased the number of possible subclusters for the FlowSOM analysis. Those changes led to the identification of 21 CD8^+^ T cell subpopulations (Figure S6B). All clusters were categorized according to their major T cell subpopulations, as previously described.

In a linear mixed model (LMM), we aimed to explore how AD biomarker response variables such as ‘Cortical β-amyloid PET’ and ‘Plasma p-tau181 levels’ were associated with covariates such as ‘Relative cell cluster abundance at each time point considered’ (=relAbundance), ‘Group’ (Accumulators versus Non-accumulators), and ‘Age’ (Formula S1) **(Singer and Willett, 2003)**. As a readout, we used an interaction term describing the combined effect of changes in relative abundance of immune cell clusters and assignment to the Accumulator group (= relAbundance:Accumulator). Using this interaction term, we investigated whether the individual AD biomarker increase over time was significantly associated with changes in relative cell cluster abundance of Accumulators, or only with natural aging (Figure 6A).

**Figure 6.**
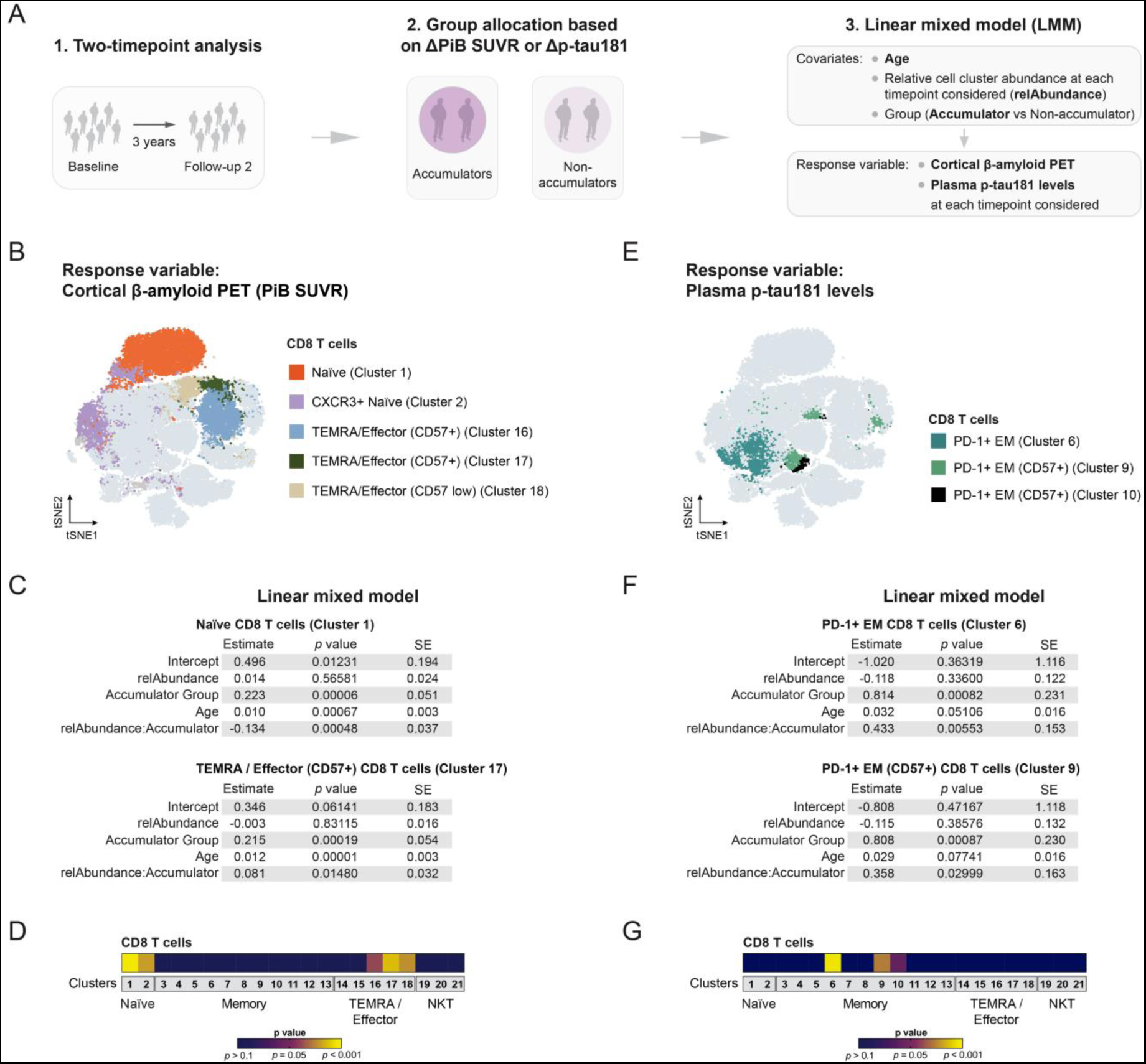
Longitudinal Analysis Indicates Specific T cell Alterations in AD Biomarker Accumulators. **(A)** Overview of the analysis strategy for the longitudinal data. ***1)*** Two-time point analysis. ***2)*** Group allocation into β-amyloid (PiB SUVR) or plasma p-tau181 Non-accumulators/Accumulators. ***3)*** Linear mixed model (LMM) with predefined response variables and covariates. The interaction term describing the combined effect of changes in relative abundance of immune cell clusters and assignment to the Accumulator group (= relAbundance:Accumulator) serves as readout. **(B-D)** LMM analysis for response variable ‘Cortical β-amyloid PET (PiB SUVR)’. CD8^+^ T cell subclusters with significant association of the interaction term relAbundance:Accumulator with PiB SUVR increase over time are highlighted in tSNE/FlowSOM map **(B)**. Detailed estimates, *p* values and standard errors (SE) are listed for the most significant associations **(C)**. A heatmap summarizes the *p* values for the association of interaction term relAbundance:Accumulator with PiB SUVR at each time point considered **(D)**. **(E-G)** LMM analysis for response variable ‘Plasma p-tau181 levels’. CD8^+^ T cell subclusters with significant association of relAbundance:Accumulator with plasma p-tau181 increase over time are highlighted in tSNE/FlowSOM map **(E)**. Detailed estimates, *p* values and standard errors (SE) are listed for the most significant associations **(F)**. A heatmap summarizes the *p* values for the association of relAbundance:Accumulator with plasma p-tau181 levels at each time point considered **(G)**.

When focusing on the response variable ‘Cortical β-amyloid PET’, we found significantly associated interaction terms for CD8^+^ T cell clusters 16, 17 and 18, all belonging to the subcluster category of TEMRA/Effector cells with CD27^−^ marker expression. The most significant association was observed for CD57^+^ TEMRA/Effector cells (CD8^+^ T cell cluster 17, relAbundance:Accumulator *p* value = 0.01480, Figure 6B-6D). In line with our previous findings, significant associations were also found for naïve CD8^+^ T cells (CD8^+^ T cell cluster 1 and 2, Figure 6B-6D).

For the response variable ‘Plasma p-tau181 levels’, we found significant associations for CD8^+^ T cell clusters 6, 9 and 10, which all belong to PD-1^+^, potentially exhausted EM T cells (Figure 6E-6G). These associations were consistent with the results of *Cross-sectional I* (Figure 2K). Clusters 6, 9 and 10 were further characterized by low CD127 and high KLRG1 expression. CD57 expression was detected for clusters 9 and 10 (Figure S6B).

In summary, the longitudinal cohort analysis also confirmed adaptive immune cell signatures that are associated with the accumulation of AD biomarkers such as brain β-amyloid deposition and increase in plasma p-tau181 levels. Importantly, these immune changes were observed during an early, preclinical stage of AD, when cognitive impairment is usually not yet present. The majority of study participants remained cognitively healthy, with the exception of two individuals who converted to MCI.

## DISCUSSION

This work demonstrates that pathological alterations in AD biomarkers such as brain β-amyloid deposition revealed by amyloid PET imaging and increases in plasma p-tau181 levels during early stages of AD pathology are associated with systemic changes within the T cell compartments of the adaptive immune system. The main strengths of our study are (I) the combined association of peripheral T cell alterations with AD biomarkers and cognitive status, (II) the consideration of both cross-sectional differences and longitudinal accumulation of cerebral β-amyloid and plasma p-tau181, and (III) the detailed analysis of latent viral background infections.

In cross-sectional analysis setups, T cells showed comparable trends in cognitively healthy β-amyloid+ individuals. We observed reduced relative abundances of naïve T cells and increased relative abundances of differentiated effector subtypes such as highly reactive CD8^+^ CD57^+^ TEMRA cells. These observations suggest an adaptive immune activation event with a shift from non-activated, naïve T cells to cells that have undergone initial antigenic stimulation and are in a stage of replicative senescence or present as reactivatable memory cells. While no consistent cell type has been reported so far, the general consensus of many previous studies has been that functional non-naïve T cell subpopulations such as memory T cell phenotypes **(Tan et al., 2002) (Larbi et al., 2009) (Pellicano et al., 2012)**, activated CD8^+^ HLA-DR^+^ T cells **(Lueg et al., 2015)** or CD8^+^ TEMRA cells **(Gate et al., 2020)** are increased in the blood of AD patients. However, previous studies mostly lack a detailed analysis of β-amyloid status and/or plasma AD biomarkers. Our results add to the existing body of evidence the important information that adaptive immune changes are in fact associated with cerebral β-amyloid status and its change over time, rather than to clinical diagnosis alone. For instance, the increased number of CD8^+^ TEMRA cells in AD has previously been interpreted as specific to the cognitive states of MCI and AD **(Gate et al., 2020)**. However, we show that elevated numbers of CD8^+^ TEMRA cells can be observed as early as in the stage of preclinical AD (HCS β-amyloid+), in which cognitively healthy subjects already present with substantial cerebral β-amyloid depositions and altered plasma AD biomarker levels. Therefore, it is essential to analyze cohorts across the entire AD continuum. As for other neurodegenerative diseases, specific T cell responses have also been associated with amyotrophic lateral sclerosis (ALS) **(Campisi et al., 2022)** and early, preclinical stages of Parkinson’s disease **(Lindestam Arlehamn et al., 2020)**. Importantly, our findings on CD8^+^ TEMRA/Effector cells can be reproduced in a setup, in which diagnostic groups are assigned based on cognitive state and levels of plasma AD biomarker p-tau181. Abnormalities in p-tau181 levels are specific for AD-related neuropathological change and not affected by potential co-morbidities such as Lewy body disease (LBD) or limbic age-related TDP-43 encephalopathy **(Smirnov et al., 2022)**. It should also be noted that previous studies that found only minimal **(Yasuno et al., 2017)** or no changes **(Stowe et al., 2017)** in peripheral immune cell populations in β-amyloid+ subjects used smaller cohort sizes and conventional flow cytometry with antibody panels unable to delineate implicated cell subtypes. We believe that our approach, based on multiparameter mass cytometry, has allowed us to preserve the complexity of cellular markers and to obtain more information from comprehensively characterized subject cohorts.

Our cross-sectional results suggest that cerebral β-amyloid accumulation is associated with specific CD8^+^ T cell alterations. However, such changes could also be caused by latent reactivated viral infections, as is suspected in AD **(Ball, 1982)**. Correlative studies have shown increased DNA content from *Herpesviridae* such as Herpes simplex virus 1 (HSV-1) in *post-mortem* brains of AD patients, specifically within β-amyloid plaques **(Jamieson et al., 1991) (Itzhaki et al., 1997) (Wozniak et al., 2009)**. Reactivation or recent infections with other *Herpesviridae* such as HSV-2, CMV and EBV have also been associated with AD development to a lesser extent **(Carbone et al., 2014) (Lovheim et al., 2018) (Warren-Gash et al., 2019)**. A different hypothesis postulates that Aβ serves as an antimicrobial peptide and aggregates as part of an innate immune response in order to provide an antiviral, protective function **(Eimer et al., 2018)**. Here, we found that preclinical β-amyloidosis-related alterations of CD8^+^ T cells are independent of latent, reactivated or acute *Herpesviridae* background infections. Importantly, in contrast to studies mentioned before, we did not measure brain or CSF virus load but serum antibody titers, since we were interested in the peripheral effect of anti-viral immune reactions.

The longitudinal cohort analysis suggests again that subtypes of naïve adaptive immune cell subpopulations and CD8^+^ TEMRA/Effector cells are associated with cerebral β-amyloid accumulation in cognitively healthy subjects. As expected, we found CD27^−^ CD45RA^+^ TEMRA/Effector cells showing CD57 expression across the full spectrum, including terminally differentiated CD57^+^, as in the cross-sectional analysis, and higher proliferative CD57^−^ subtypes **(Verma et al., 2017)**. In line with cross-sectional results, we also found an association of PD-1^+^ EM cells with increasing plasma p-tau181 levels. Those cells are potentially exhausted due to persistent antigenic stimulation and upregulate inhibitory receptors including Programmed Death-1 (PD-1 or CD279) **(Pauken et al., 2016)**. In *Cross-sectional I*, we observed a reduction of these PD-1^+^ EM in HCS β-amyloid+ compared to HCS β-amyloid–, probably indicating a shift of abundances from exhausted cell types with impaired effector functions to highly antigen-specific TEMRA/Effector cells. Another possibility would be that PD-1^+^ EM represent a putative precursor for TEMRA/Effector cells, since their IL-7 receptor (CD127) expression is already reduced, while some subtypes upregulate KLRG1 and CD57 expression.

Since major immune changes were observed in a preclinical stage of AD, the adaptive immune system might respond directly or indirectly to an initial change in AD biomarkers. Possible epitope targets of the identified T cells are still subject to speculation at the moment and can only be revealed by targeted epitope mapping for clonally expanded cells. As the immune system is exposed to autologous Aβ peptide long before its deposition in β-amyloid plaques, it is very likely that Aβ-specific T cells are negatively selected in the thymus during early development, or are subject to peripheral tolerance. Therefore, the native Aβ1-42 peptide itself is unlikely to elicit such an (auto-)immune response, since several tolerance barriers would have to be overcome. Moreover, CD8^+^ cytotoxic T cells are not the T cell subtype that is expected to respond to mainly extracellular threats such as β-amyloid depositions. However, there are ways how such an adaptive immune response could be initiated. For example, expanded T cell clones discovered in some neurodegenerative/neurological disorders showed specificity against viral antigens **(Luo et al., 2018) (Wang et al., 2020)**. These T cell responses could trigger bystander activations of innate antigen-presenting cells and adaptive immune responses on the one hand. On the other hand, infectious diseases might also be the trigger for the development of an autoreactive immune response through molecular mimicry between peptides derived from infectious agents and self-peptides. Such antigen cross-recognition does not require perfect homology of epitope sequences, but may occur when peptides share common amino acid motifs. Also, the concept of T cell neo-epitopes, against which there is tolerance in only a few cases, has to be further explored as potential source of TEMRA cell activation **(Rickenbach and Gericke, 2021)**.

We conclude that changes within the peripheral blood T cell compartments precede the clinical diagnosis of AD, are associated with brain β-amyloidosis and/or plasma AD biomarkers, and are independent of latent viral infections. Future research will be directed towards confirmatory longitudinal studies with larger cohort size and longer follow-up. This information could be used to further investigate whether immune cell changes might even precede established AD plasma biomarkers, as they can respond to low peptide antigen concentrations, whereas certain peptide levels above the detection limit are required for successful plasma analysis. Early AD-associated immune cell changes might provide guidance and prompt more expensive downstream analysis techniques if a population at risk is found. It has to be investigated whether the observed immune cell changes consistently disappear with progressing AD pathology as observed for the MCI β-amyloid+ subjects of *Cross-sectional I* who had the highest cerebral β-amyloid load. It might be that due to constant exposure to an antigenic stimulus, which drains from the preclinical AD brain into the periphery via cerebral lymphatics, an initial immune response is abrogated by mounting tolerance mechanisms or immunological exhaustion. In addition, it remains to be elucidated whether these initial immune responses are indeed protective against toxic protein aggregates in the brain or other antigenic entities related to AD, or whether they even exacerbate the pathology. In fact, in aging studies, cognitively healthy, potentially resilient centenarians seemed to have higher numbers of effector T cells in peripheral blood compared to younger control subjects **(Fagnoni et al., 1996) (Hashimoto et al., 2019)**. In contrast, many studies have shown a lack of proper immune responses in mouse models of β-amyloidosis and AD **(Ferretti et al., 2016) (Gericke et al., 2020) (Bettcher et al., 2021)**. Thus, it is perhaps desirable to have less exhausted immune cells such as PD-1^+^ EM cells and more antigen-specific effector T cells. To further determine in particular the importance of CD8^+^ CD57^+^ TEMRA/Effector cells in AD pathology, it would be necessary to confirm the presence of this cell type in AD brain tissue and long-term longitudinal studies.

In summary, we propose that cellular immune patterns in the periphery could be used as a proxy for early AD biomarker changes. A better understanding of these immune patterns in combination with detailed epitope specificity analysis might pave the way for the identification of new composite biomarker candidates and complement the diagnostic toolbox for the detection of AD, in particular preclinical AD. Moreover, understanding the pathophysiological significance of these processes may give rise to new therapeutic strategies.

## Supporting information

Supplemental Formula S1

## ACKNOWLEDGEMENTS

We would like to thank all the volunteers who kindly participated in the studies. We thank all study physicians and neuropsychologists for their contribution to the assessments and conduct of the study. Samples from participants were collected at the Center for Prevention and Dementia Therapy (Institute for Regenerative Medicine, University of Zurich) by study nurses led by Esmeralda Gruber. We would like to thank the Cytometry Facility (University of Zurich) for technical assistance, and the Institute of Medical Virology (University of Zurich) for viral serology analysis. This work was supported by institutional funding of the University of Zurich, grants from the Synapsis Foundation – Alzheimer Research Switzerland ARS (No. 2019-PI06 to R.M.N., C.G., and A.G.), the Swiss National Science Foundation (SNF 33CM30-124111, SNF 320030-125387/1 to C.H.), the Mäxi Foundation (to C.H.) and the Velux Foundation (to L.K.).

## AUTHOR CONTRIBUTIONS

Conceptualization, C.G., T.K., L.K., V.Treyer, M.T.F., and A.G.; Methodology, C.G., T.K., and V.Tosevski; Software, R.F., C.R., and V.Tosevski; Formal Analysis, C.G., T.K., and R.F.; Investigation, C.G., T.K., A.M., and C.R.; Resources, L.K., V.Treyer, and A.G.; Data Curation, R.F.; Writing - Original Draft, C.G. and T.K.; Writing - Review & Editing, C.G., T.K., R.F., L.K., C.H., V.Treyer, M.T.F., and A.G.; Visualization, A.M.; Supervision and Project Administration, C.H. and R.M.N.; Funding Acquisition, C.G., L.K., C.H., R.M.N., M.T.F. and A.G.

## DECLARATION OF INTERESTS

T.K. is currently an employee of Charles River Associates, Switzerland; L.K. is an employee of Roche, Switzerland; C.H. and R.M.N. are members of the board of directors and shareholders of Neurimmune AG, Switzerland; M.T.F. is co-founder and CSO of the Women’s Brain Project; M.T.F. has received in the past two years personal fees from Roche, Lundbeck, GW, for activities unrelated to this paper.

## METHODS

### RESOURCE AVAILABILITY

#### Contact for Reagent and Resource Sharing

Further information and requests for resources and reagents should be directed to and will be fulfilled by the Lead Contact, Dr. Christoph Gericke.

#### Materials Availability

This study did not generate new unique reagents.

#### Data and Code Availability

The data sets and code generated during this study are publicly available at Zenodo (CERN, European Organization for Nuclear Research) under https://doi.org/10.5281/zenodo.7157682.

### EXPERIMENTAL MODEL AND SUBJECT DETAILS

#### Ethics Statement

The participants included in the analyses participated in in-house research studies conducted by the ‘Center for Prevention and Dementia Therapy’ at the Institute for Regenerative Medicine at the University of Zurich, Switzerland. All participants gave written informed consent. All studies as well as further use for the current analyses were approved by and conducted in concordance with the guidelines of the local ethics committee (Kantonale Ethikkommission Zürich) and the Declaration of Helsinki **(World Medical Association, 2013)**.

#### Study Participants – Exploratory Cross-sectional Study I

In an exploratory study named ‘*Cross-sectional I*’, we included cryopreserved peripheral blood mononuclear cells (PBMCs), blood plasma and serum from cognitively healthy control subjects (‘HCS’, n=23), subjects diagnosed with mild-cognitive impairment (‘MCI’, n=18) according to consensus criteria **(Winblad et al., 2004)**, and patients with probable Alzheimer’s disease dementia (‘AD’, n=9) according to NINCDS-ADRDA 1980 **(McKhann et al., 1984)**, ICD-10 criteria **(World Health Organization, 1993)**. All subjects previously participated in longitudinal cohort studies with comparable research protocols. Cerebral β-amyloid load was assessed by positron emission tomography (PET) imaging with [11C]-Pittsburgh compound B (PiB) tracer except for AD subjects who were assessed in a protocol without β-amyloid-PET. We used the standardized uptake value ratios (SUVR) and the Centiloid method for better comparability among different tracers **(Klunk et al., 2015)**. Subjects were selected to compare low PiB SUVR (β-amyloid negative) and high PiB SUVR clearly above the cut-off (β-amyloid positive). The lowest PiB+ SUVR was 1.514 while the highest PiB-SUVR was 1.250, the pre-defined SUVR cut-off level was 1.265 **(Gietl et al., 2015)**. Study participants received extensive blood examinations including apolipoprotein E genotyping (*APOE ε2, ε3, ε4)*, viral antibody titer analysis (against Herpesviruses) and assessment of acute inflammatory markers (C-reactive protein, CRP). The median time difference between PBMC sampling and PET imaging was 19 days. MCI β-amyloid– subjects were included as internal control, since they are considered to be cognitively impaired due to reasons other than AD. MCI β-amyloid+ patients showed both early signs of cognitive impairment and β-amyloid deposition, and were categorized as ‘MCI due to AD’. Detailed group sizes, demographics and characterization of *Cross-sectional I* are shown in Figure 1C, Figure S1 and Table S1. Due to the exploratory nature of *Cross-sectional I*, no *a priori* power analysis was conducted to estimate sample sizes.

#### Study Participants – Validation Cross-sectional Study II

A second cross-sectional study, termed ‘*Cross-sectional II*’, included 95 HCS and 47 MCI cases from the baseline of an ongoing longitudinal cohort study whose participants were recruited via newspaper advertisements. Neuropsychological examination according to consensus criteria and further blood examinations were performed in a comparable way as in *Cross-sectional I*. All subjects received PET imaging with [18F]-Flutemetamol (FMM) tracer (Vizamyl, GE Healthcare). Finding optimal cut-offs was important for the cohort design, therefore we included FMM Centiloids representing ‘AD biomarker-positive’ cases (12 < Centiloid < 30) and ‘established AD pathology’ cases (Centiloid > 30) **(Salvado et al., 2019)**, with an overall cut-off of FMM Centiloid > 12 for β-amyloid+ subjects. All β-amyloid– cases had Centiloid levels below 9. This resulted in a more homogenous and therefore more natural β-amyloid PET signal distribution compared to *Cross-sectional I,* including β-amyloid+ subjects with FMM SUVR and Centiloid close to the cut-off (Figure S1H, S1I and S1K). Further selection was based on health status (excluding current/recent tumor patients or patients with Hashimoto’s disease) and age range. Considering these criteria, *Cross-sectional II* contained 50 HCS β-amyloid–, 45 HCS β-amyloid+, 26 MCI HCS β-amyloid– and 21 MCI HCS β-amyloid+ subjects. According to *a priori* power analysis considering the effect size in CMV titer-adjusted CD8^+^ TEMRA/Effector cell analysis (Figure 4D), the minimum sample size for a validation HCS β-amyloid+ group should be n = 26 (Power = 0.8; Type I error = 0.1) **(Glueck, 2008)**. Detailed group sizes, demographics and characterization of *Cross-sectional II* are shown in Figure 1D, Figure S1 and Table S2.

#### Study Participants – Longitudinal Study

For the longitudinal study, participants were followed over a total time period of 3 years. In order to be eligible, subjects had to be between 55 and 80 years old and cognitively healthy at baseline which was ascertained by an MMSE (mini-mental state examination) minimum score of 27 **(Folstein et al., 1975) (Tombaugh and McIntyre, 1992)** as well as comprehensive clinical and neuropsychological examination. Every eligible subject for the current analysis (n=59) received two separate [11C]-PiB-PET scans at *Baseline* and after 3 years at the end of the study. PBMCs were isolated from participants’ blood donations at *Baseline*, after 18 months (=’*Follow-up1*’) and after 3 years (=‘*Follow-up2*’). While all individuals received β-amyloid PET imaging at *Baseline* and *Follow-up2*, PBMC samples were not available for all visits of each subject. Availability of PBMCs from different visits is summarized in Table S3. Due to the exploratory nature of this longitudinal study, no *a priori* power analysis was conducted to estimate sample sizes.

In the longitudinal study, subjects were considered β-amyloid+ at PiB SUVR > 1.223 (75th percentile of baseline SUVR). We allocated subjects into groups of cerebral β-amyloid ‘Non-accumulators’ and β-amyloid ‘Accumulators’ based on their change in cortical PiB SUVR over time (termed ‘ΔPiB’). Previous studies have used basic discriminations considering every subject with an annual ΔPiB SUVR > 0 as Accumulator **(Villemagne et al., 2013) (Landau et al., 2018)** or have established cut-offs based on inflexion points between bimodal distributions of ΔPiB SUVR/year **(Villain et al., 2012)**. Because negative ΔPiB SUVR in *Baseline* β-amyloid– subjects presumably reflects measurement noise **(Landau et al., 2018)**, we allocated individuals with ΔPiB SUVR < 0 automatically to a group of Non-accumulators. We aimed to identify only Accumulators with clearly increasing PiB retention over time compared to Non-accumulators. For that reason, all individuals with ΔPiB SUVR ≥ 0 were split by median, and as a result subjects with ΔPiB SUVR ≥ 0.04 were classified as Accumulators, whereas subjects with ΔPiB SUVR < 0.04 were categorized as Non-accumulators. In an alternative group allocation, we divided subjects according to the change of plasma p-tau181 levels over time (Δp-tau181). Via median split, we defined a cut-off at Δp-tau181 = 0.13pg/ml, to distinguish between p-tau181 Accumulators and Non-accumulators. Detailed group sizes, demographics and characterization of the cohort are shown in Figure 5D, Figure S5 and Table S4.

### METHOD DETAILS

#### APOE Genotyping

*APOE* genotyping was either performed by restriction isotyping as previously described **(Hixson and Vernier, 1990)** or by commercially available Sanger Sequencing (Microsynth AG).

#### PET Imaging and Image Analysis

Dynamic [11C]-PiB-PET or [18F]-FMM-PET images were acquired over 70 minutes on whole body PET/CT cameras in 3D mode (Discovery RX and Discovery STE, GE Healthcare). Subjects received approximately 350 MBq of tracer. Image reconstruction included standard corrections and filtered back projection algorithms with an image resolution of 2.34*2.34*3.27 mm. For magnetic resonance imaging (MRI), 3D TFE T1-weighted imaging was performed on a Philips 3T Achieva with an image resolution of 0.9375*0.9375*1 mm.

Images were processed with PMOD 3.7 NeuroTool (PMOD Technologies, Zurich, Switzerland) and brain parcellation method. MR images were segmented (‘3 probability maps methods’) and cortical regions of interest were defined according to ‘Adult brain maximum probability map’ by Hammers et al. as implemented in PMOD NeuroTool. For non-cortical region definitions, deep nuclei parcellation method as implemented in NeuroTool was performed (each hemisphere is processed individually, 16 reference sets were taken into account to improve accuracy). Grey matter–white matter segmentation cut-off was defined at a standard 50% probability. This segmentation was applied to region of interest definitions to exclude white matter voxels from analysis. For the follow-up scans in the longitudinal study, the same MR baseline image and the same regions of interest definitions were taken to reduce variability due to automatic segmentation and outlining process between the two time points. PET images were co-registered to the baseline MR images based on the early frames signal (6 min) of the dynamic PET scan, which resembles a perfusion map. Manual correction was applied where needed to ensure optimal co-registration. On the fused individual images in MR space, correct outlining of the regions of interest was checked and corrected if necessary. Late phase uptake values were calculated as average late frames activity (50 to 70 minutes post injection) of cortical regions (averaging cortical regions as defined by the ‘Centiloid’ approach and divided by cerebellar cortex activity) **(Klunk et al., 2015)**. Due to the standard procedure and standard calibration of equipment, the SUVR approach was considered sufficient and most robust to evaluate changes over three years in the longitudinal study cohort.

#### Neuropsychological Testing

The test selection for the descriptive analysis was based on availability of identical neuropsychology tests over all cohorts and should provide information on the cognitive status or changes of the different subgroups. In all studies, the participants received first a short informal interview to assess subjective cognitive complaints and factors with possible influence on the test results (e.g. alcohol/drug abuse, medication, recent travelling over time zones) as well as characteristics of free language expression. The here reported neuropsychological tests (Table S5) were comparable in all studies but were not necessarily performed in the same order. Especially in the AD group, not all available tests were performed. Z-scores (not reported, available upon request) were calculated based on norm values of the individual tests as defined in the respective test manuals.

#### Plasma Biomarker Analysis

Blood plasma samples were obtained on the same day as PBMC extraction was performed. Blood of study participants was collected in EDTA tubes (Vacutainer® EDTA Tubes, BD), inverted 10 times and centrifuged at 1620 x g for 12min at 6°C. Plasma supernatant was aliquoted and immediately stored at -80°C. For ultrasensitive AD biomarker quantification, EDTA-plasma of Cross-sectional I and the longitudinal study was shipped to Quanterix corporation (Quanterix, Billerica, MA, USA). Analysis was performed on a Simoa HD-X instrument using Simoa Human Neurology 3-plex A (N3PA) and P-tau181 V2 immunoassays for measuring plasma Aβ1-42, Aβ1-40, total tau and phospho-tau181 concentrations.

#### CyTOF Antibody Panels

Monoclonal, heavy metal isotope-conjugated anti-human antibodies were purchased from Fluidigm or self-made from purified antibodies and heavy metal isotopes using the Maxpar X8 Multi-Metal Labeling Kit (Fluidigm) according to manufacturer’s instructions. Antibody panels were designed for barcoding purposes as well as for characterization of T cells for both cross-sectional and longitudinal studies. All panels and further panel information are listed in the Supplemental Materials.

#### CyTOF Barcoding Strategies

In order to reduce technical noise due to staining variabilities among single samples, we applied different barcoding strategies to allow for sample pooling and bulk PBMC analysis. For all studies we applied custom-made live-cell barcoding based on combinations of anti-human CD45 antibodies labelled with different metal tags. For the T cell study in *Cross-sectional I*, we applied an 8-choose-3 approach (56 possible combinations), whereas for *Cross-sectional II* and the longitudinal T cell study we applied a 10-choose-4 approach (210 possible combinations). Detailed live-cell barcoding panels are listed in Table S6.

#### Sample Preparation and Surface Staining

Cryopreserved PBMCs were thawed in a 37°C water bath and transferred into tubes with pre-warmed cell medium (RPMI 1640, Sigma Aldrich with 10% heat-inactivated FBS (Gibco) and 1% Glutamax (Gibco)). After two washes in cell medium (350 x g, 7min), 1 million PBMCs from each subject were transferred into a 96-well plate, washed twice in PBS (350 x g, 5min) and incubated with Cell-ID Cisplatin-^198^Pt (Fluidigm) 1:10’000 in PBS for dead-cell exclusion staining for 10min at room temperature (RT). Cells were incubated with Fc receptor blocking antibodies (Human TruStain FcX, Biolegend) 1:20 in Cell Staining Buffer (CSB, Fluidigm) for 5 minutes at RT before applying the respective barcoding strategies. After barcoding, cells were washed twice in CSB, pooled and incubated as bulk sample with their antibody cocktail (see Supplemental Materials) for 20 minutes at RT. After washing in CSB, cells were fixed with 1.6% formaldehyde in CSB and stored overnight at 4°C. The next day, stained samples were aliquoted in max. 20 Mio. cell batches and frozen in 10% dimethyl sulfoxide (DMSO, Thermo Fisher) in FBS for long-term storage at -80°C. On day of analysis, frozen samples were thawed in pre-warmed PBS and acquired on the mass cytometer (CyTOF) according to acquisition protocol (Figure 1B).

#### CyTOF Acquisition

Prior acquisition, cells were incubated in Cell-ID Intercalator-Ir (Fluidigm) 1:5000 in Maxpar Fix and Perm Buffer (Fluidigm) for 1 hour at RT. After one wash step in deionized metal-free water and two wash steps in Cell Acquisition Solution (CAS, Fluidigm), samples were diluted to 1.2 million cells/ml in CAS containing 1:10 EQ Four Element Calibration Beads (Fluidigm). Samples were analyzed with a Helios CyTOF2 mass cytometer (DVS Sciences, Fluidigm).

#### Serology for Virus Antibody Titers

Viral antibody titers in serum samples from both cross-sectional studies were analyzed via ELISA or indirect immunofluorescence at the Institute of Medical Virology, University of Zurich. Antibody titers against Cytomegalovirus (CMV) (IgG and IgM), Epstein-Barr virus (EBV) (IgG and IgM against viral capsid antigen (VCA) and IgG against Epstein-Barr nuclear antigen (EBNA) were tested), Herpes simplex virus (HSV) (IgM for HSV-1+2, HSV-1 IgG, HSV-2 IgG) and Varicella-Zoster-Virus (VZV) (IgG and IgM) were investigated. Quantitative results were obtained for the following antibody titers: VZV IgG (in mIU/ml, threshold for positive test >100), CMV IgG (in AE/ml, threshold ≥6), HSV-1 IgG (in ‘Signal divided by cut-off’: S/CO, threshold >1) and HSV-2 IgG (in S/CO, threshold >1). All other titers were analyzed qualitatively (positive/negative).

#### C-reactive Protein Assessment

C-reactive protein (CRP) levels for serum samples from both cross-sectional and longitudinal study participants were analyzed as part of routine diagnostics externally in certified laboratories to exclude acute background infections and inflammation.

## QUANTIFICATION AND STATISTICAL ANALYSIS

### CyTOF Data Preprocessing

Bead normalization over time, file concatenation, debarcoding and compensation for overspilling high-intensity barcoding channels were performed using Premessa (R package, version 0.2.6) and Catalyst (R package, v1.14.0) in an R environment (R, v4.0.2) implemented in RStudio (v1.3.1093). All R packages used are listed in the Supplemental Materials.

### Multivariate Data Analysis

Custom-made R scripts (see data/code repository) were used for data analysis and visualization. Mass signals from calibration beads, cisplatin-positive (^198^Pt) dead cells, iridium-low (^191^Ir, ^193^Ir) debris and iridium-high (^191^Ir, ^193^Ir) cell doublets were cleaned out. For further analysis, CD45^+^ PBMCs, CD8^+^ T cells (defined as CD3^+^ CD4^−^ CD8^+^) and CD4^+^ T cells (defined as CD3^+^ CD4^+^ CD8^−^) were selected and exported as individual cell subsets. A maximum of 10’000 events (CD45+ leukocyte analysis and T cell analysis) per FCS file were randomly selected and processed via flowCore (R package, v2.0.1). In case of files that included less than 10’000 events, all events were used for the analysis. All events were transformed using an inverse hyperbolic sine (arcsinh) function with a cofactor of 5 **(Nowicka et al., 2017)**. Next, the multivariate n-dimensional data was processed by tSNE (*t*-distributed stochastic neighbor embedding), a nonlinear dimensionality reduction algorithm (Rtsne package, v0.15). Spatial separation of events in the resulting tSNE map is controlled by the ‘perplexity’ parameter.

### FlowSOM Clustering

Automated and unsupervised clustering was performed in R using the FlowSOM algorithm **(Van Gassen et al., 2015)** implemented in the FlowSOM R package (v1.20.0). For meta-clustering, ‘consensus hierarchical clustering’ was used and respective k-values defining the number of resulting clusters were determined based on estimations of expectable subclusters **(Hartmann et al., 2016)**.

### Identification of Immune Cell Subsets

Via data visualization package ggplot2 (R package, v3.3.2) automatically identified FlowSOM clusters were overlaid with their corresponding cell events in the tSNE map, and heatmaps displaying median signal intensities per cluster were created. Based on signal intensities, immune cell subpopulations were identified and similar clusters were merged to reflect biologically meaningful immune cell subsets. Eventually, relative cluster abundances were determined as quantification readout.

### CITRUS

In parallel to FlowSOM, CITRUS (Cluster Identification, Characterization and Regression) was used as an alternative method for unsupervised clustering and identification of statistically significant differences in immune cell abundances between diagnostic groups **(Bruggner et al., 2014)**. The correlative significance analysis of microarrays (SAM) association model was used with maximum 10’000 cells per sample, 1% minimum cluster size and 5% false discovery rate (FDR). Clusters that showed differential relative abundances among diagnostic groups were quantified and visualized.

### Cross-sectional Studies – Statistical Analysis

Statistical analysis was performed using Graphpad Prism (v9.1.2) and R (v4.0.2). Assumption of normal distribution of variables was tested with the Shapiro-Wilk normality test (α = 0.05). If the null-hypothesis of normality was not rejected, significant differences between two groups were tested with a two-tailed, unpaired Student’s t test. Correlations for normally distributed data were calculated with Pearson’s correlation (α = 0.05, confidence interval 95 %, two-tailed). In case of a deviation from normal distribution, significant differences between two groups were tested with a non-parametric, two-tailed, unpaired Mann-Whitney test. Correlations were calculated with non-parametric Spearman’s correlation (α = 0.05, confidence interval 95 %, two-tailed). The difference of cluster abundances between multiple diagnostic groups was compared using the non-parametric Kruskal-Wallis test. All *p* values were corrected for multiple comparisons within each cluster using the Benjamini-Hochberg false discovery rate (FDR) method. Age-, sex- or virus-related effects were estimated and controlled using linear regression models, and analysis was performed on the basis of the corresponding residuals. All *p* values are reported as absolute values including an estimate of FDR in each figure. Due to small cohort size, significance was defined as FDR < 10%.

### Longitudinal Study – Descriptive Analysis and Statistical Modelling

For the longitudinal study, a linear mixed model (LMM) was fitted in R. For simplicity, a two-time point analysis was chosen. If available, the relative abundance of immune cell clusters and the corresponding cortical PiB SUVR or plasma p-tau181 data at time points *Baseline* and *Follow-up2* were used. If there was no immune cell data available at *Baseline* or *Follow-up2*, time point *Follow-up1* served as replacement. Cortical PiB SUVR or plasma p-tau181 at each time point considered was used as response variable (see ‘response vector’ in Formula S1). Cortical PiB SUVR and plasma p-tau181 were measured only at *Baseline* and *Follow-up2*, hypothetical values for *Follow-up1* were generated by linear interpolation (e.g. imputed PiB SUVR values in Figure S5B). The relative abundance of immune cell clusters at each time point considered was used as covariate. The LMM was used to further investigate the association of individual changes in cortical PiB SUVR or plasma p-tau181 over time with covariates such as relative abundance of immune cell clusters, group allocation (PiB SUVR or plasma p-tau181 Accumulators versus Non-accumulators) and age (see ‘design matrix for fixed effects’ in Formula S1). Additionally, we included an interaction term describing the combined effect of changes in relative abundance of immune cell clusters over tim and allocation to the Accumulator group (see ‘design matrix for fixed effects’ in Formula S1). Relative cell abundances were transformed using the arcsinh transformation. The LMM was fitted with the lme4 R package (v1.1-26).

## SUPPLEMENTAL FIGURES

**Figure S1.**
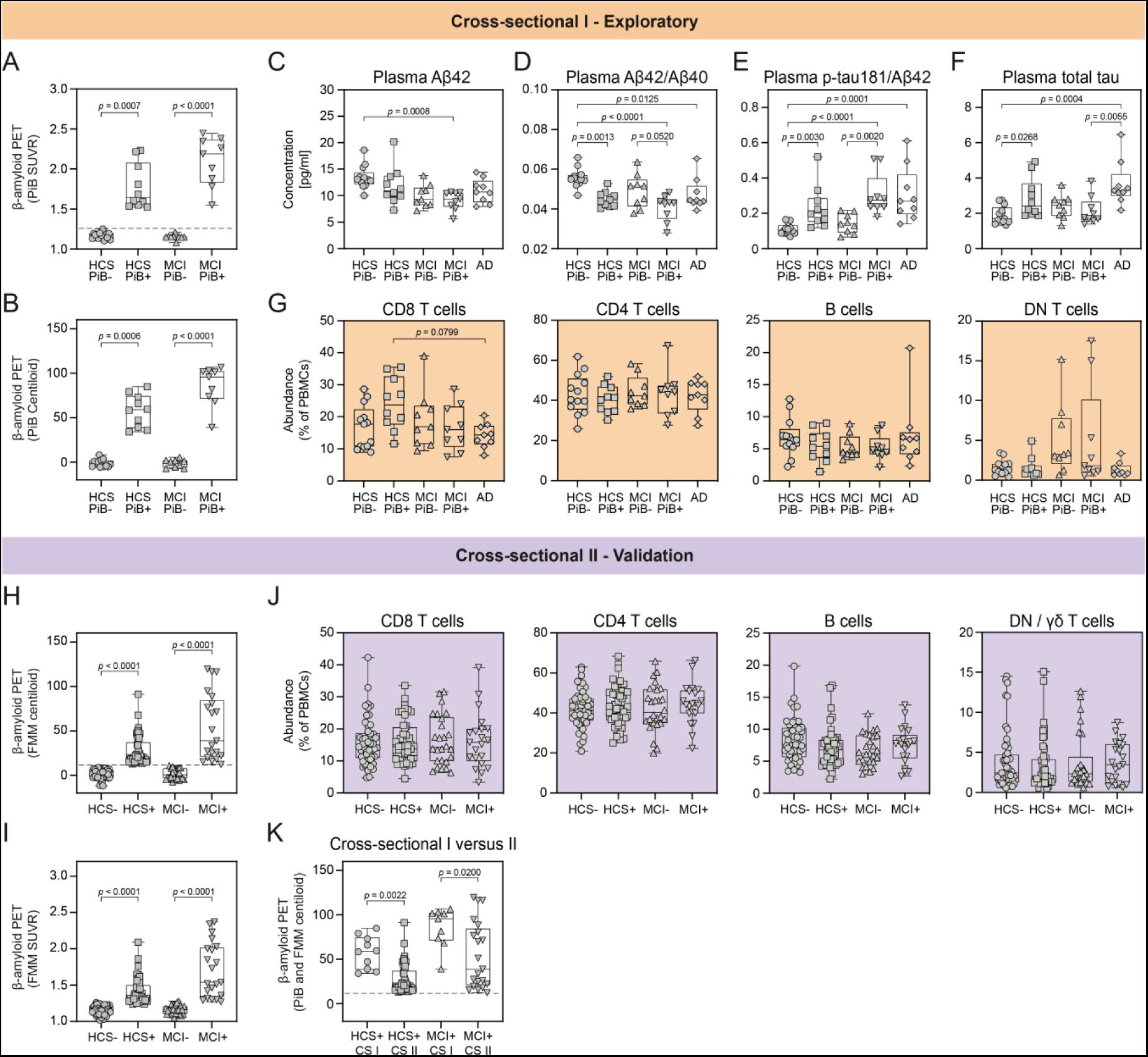
Cross-sectional Studies: General Characterization. *Cross-sectional I*: **(A, B)** Comparison of cerebral β-amyloid load between diagnostic groups in units of PiB SUVR **(A)** and PiB Centiloid **(B)**. β-amyloid+ (=PiB+) groups are significantly different from β-amyloid– (=PiB–) groups. Dashed line represents PiB SUVR cut-off for β-amyloid+ subjects (PiB SUVR = 1.265). **(C)** Plasma Aβ42 quantification shows decreased levels in MCI PiB+ compared to HCS PiB–. **(D)** Ratio of plasma Aβ42/Aβ40 is reduced in HCS PiB+ vs. HCS PiB–, MCI PiB+ vs. MCI PiB–, MCI PiB+ vs. HCS PiB– and AD vs. HCS PiB– **(E)** Ratio of plasma p-tau181/Aβ42 is increased in HCS PiB+ vs. HCS PiB–, MCI PiB+ vs. MCI PiB–, MCI PiB+ vs. HCS PiB– and AD vs. HCS PiB–. **(F)** Plasma total tau quantification shows increased levels in HCS PiB+ vs. HCS PiB–, AD vs. HCS PiB– and AD vs. MCI PiB+. **(G)** Quantification and comparison of major adaptive immune cell clusters such as CD8^+^ and CD4^+^ T cells, B cells, double-negative (DN) T cells. Differences have been observed for CD8^+^ T cells. *Cross-sectional II*: **(H, I)** Comparison of cerebral β-amyloid load between diagnostic groups in units of FMM Centiloid **(H)** and FMM SUVR **(I).** β-amyloid+ groups are significantly different from β-amyloid– groups. Dashed line represents FMM Centiloid cut-off for β-amyloid+ subjects (FMM Centiloid = 12). **(J)** Quantification and comparison of major adaptive immune cell clusters such as CD8^+^ and CD4^+^ T cells, B cells, DN / γδ T cells. No differences have been observed as previously in *Cross-sectional I*, probably due to the higher number of subjects in *Cross-sectional II* and the more homogeneous distribution of cerebral β-amyloid load. **(K)** Comparison of cerebral β-amyloid load (in PiB / FMM Centiloid) within the β-amyloid+ groups of *Cross-sectional I* and *Cross-sectional II*. PiB Centiloid in HCS β-amyloid+ and MCI β-amyloid+ of *Cross-sectional I* is significantly higher than FMM Centiloid in HCS β-amyloid+ and MCI β-amyloid+ of *Cross-sectional II*. Each symbol in quantification graphs represents data from one subject. Box plot represents interquartile range (IQR), horizontal line indicates median, whiskers extend to min/max data point. The *p* values were calculated via nonparametric Kruskal-Wallis tests. Controlling for multiple comparisons was performed with FDR method of Benjamini and Hochberg. Selected *p* values ≤ 0.1 (FDR 10%) are displayed.

**Figure S2.**
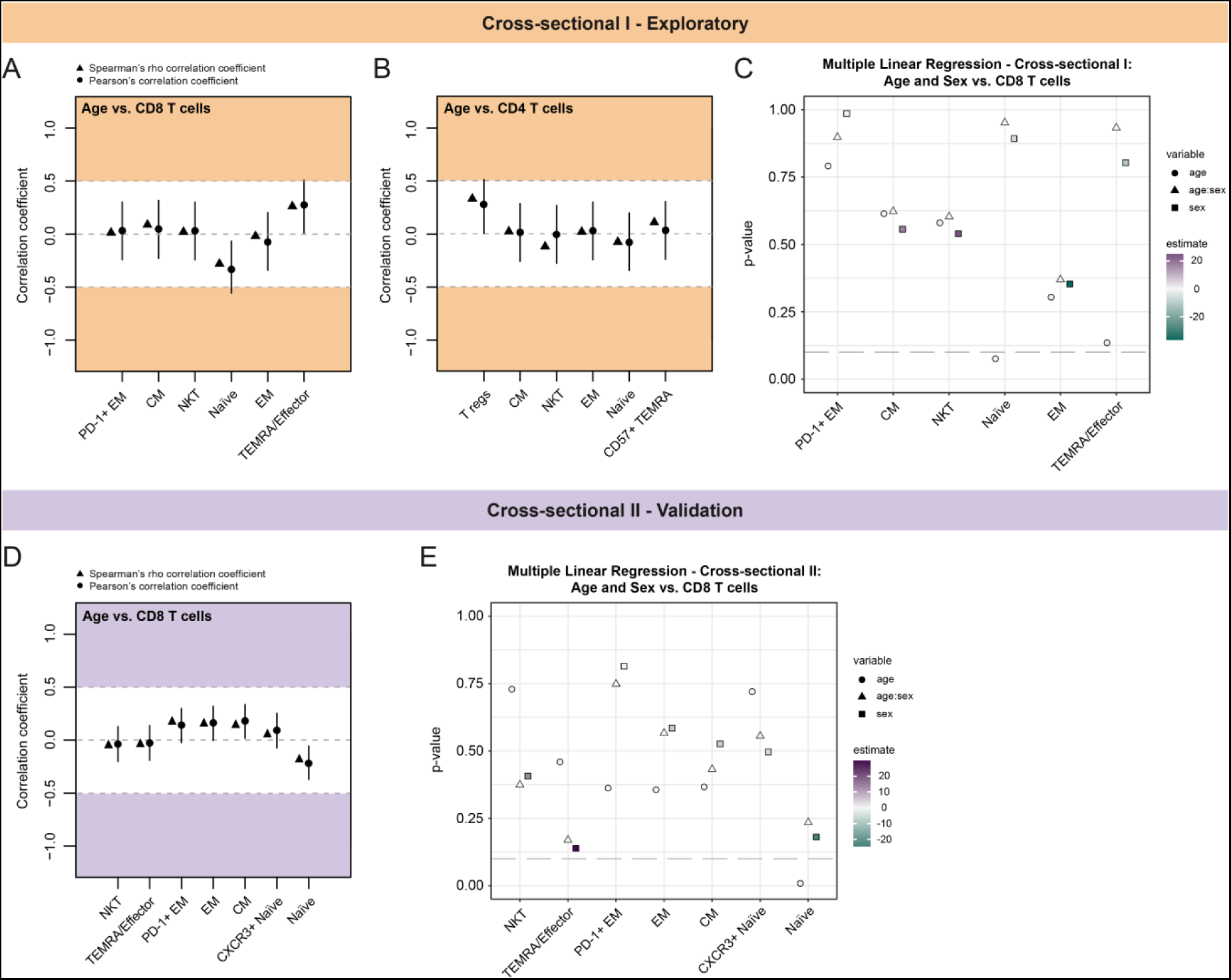
Cross-sectional Studies: Age Correlations. *Cross-sectional I*: **(A)** Correlations between ages of all subjects and relative CD8^+^ T cell cluster abundance. No substantial differences have been found. **(B)** Correlations between ages of all subjects and relative CD4^+^ T cell cluster abundance. No substantial differences have been found. **(C)** Multiple linear regression for analysis of the influence of age and sex on CD8^+^ T cell cluster abundance. No effect on CD8^+^ T cell subclusters was found, except for naïve CD8+ T cells, which were significantly affected by age, but only with a low estimate close to 0. *Cross-sectional II*: **(D)** Correlations between ages of all subjects and relative CD8^+^ T cell cluster abundance. No substantial differences have been found. **(E)** Multiple linear regression for analysis of the influence of age and sex on CD8^+^ T cell cluster abundance. No effect on CD8^+^ T cell subclusters was found, except for naïve CD8+ T cells, which were significantly affected by age, but only with a low estimate close to 0. Correlations: Black dots represent the estimated Pearson’s r correlation coefficients and the black lines the corresponding 95% confidence intervals. Black triangles represent the estimated Spearman’s rho correlation coefficients. Dashed lines are indicating a correlation cut-off at rho = 0.5 or rho = -0.5. Multiple linear regression: The *p* values for the age and sex variables and the age:sex interaction term in each CD8^+^ T cell subcluster are plotted. All data points are colored according to estimate of effect. Grey dashed line indicates *p* value of 0.1.

**Figure S3.**
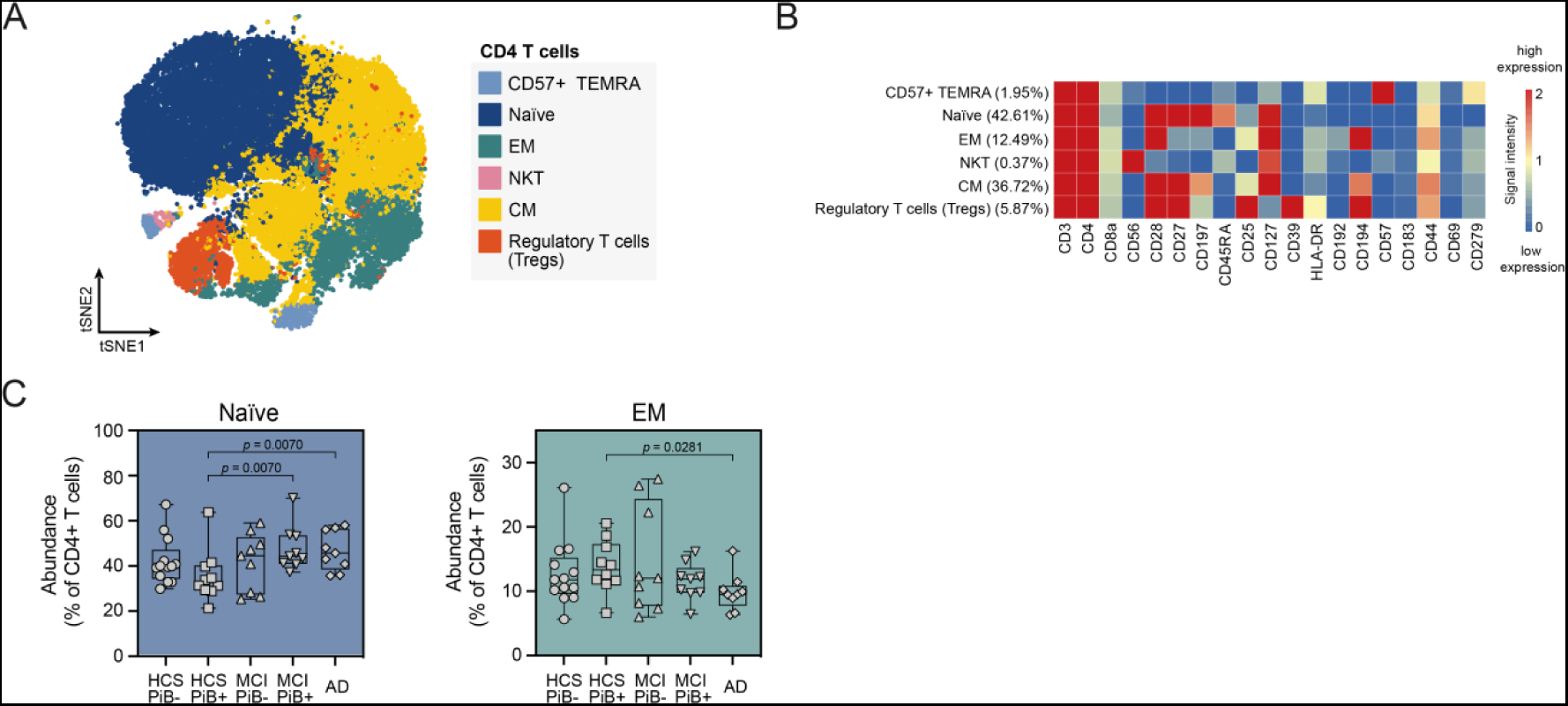
*Cross-sectional I* - Healthy β-amyloid+ Subjects Are Further Characterized by Altered CD4^+^ T cell Subpopulations. **(A)** tSNE and FlowSOM clustering for PBMC-derived CD4^+^ T cells averaged over all subjects of *Cross-sectional I* (tSNE settings: iterations = 2’000, events = max. 10’000, perplexity = 50; FlowSOM settings: k17, merged into 6 major clusters). EM = effector memory cells, NKT = natural killer T cells, CM = central memory cells. **(B)** Heatmap for identification of 6 major CD4^+^ T cell clusters. Heatmap shows *arcsinh*-transformed median marker intensities. **(C)** Quantification and comparison of relative abundances of CD4^+^ T cell subpopulations in HCS PiB+ subjects with HCS PiB–, MCI PiB+ and AD patients. Naïve CD4^+^ T cells show reduced abundance in HCS PiB+ individuals compared to MCI PiB+ and AD groups. In contrast, a higher relative abundance of CD4^+^ EM T cells has been observed in HCS PiB+ subjects compared to AD patients. Each symbol in quantification graphs represents data from one subject. Box plot represents IQR, horizontal line indicates median, whiskers extend to min/max data point. The *p* values were calculated via nonparametric Kruskal-Wallis tests. Controlling for multiple comparisons was performed with FDR method of Benjamini and Hochberg. Selected *p* values ≤ 0.1 (FDR 10%) are displayed.

**Figure S4.**
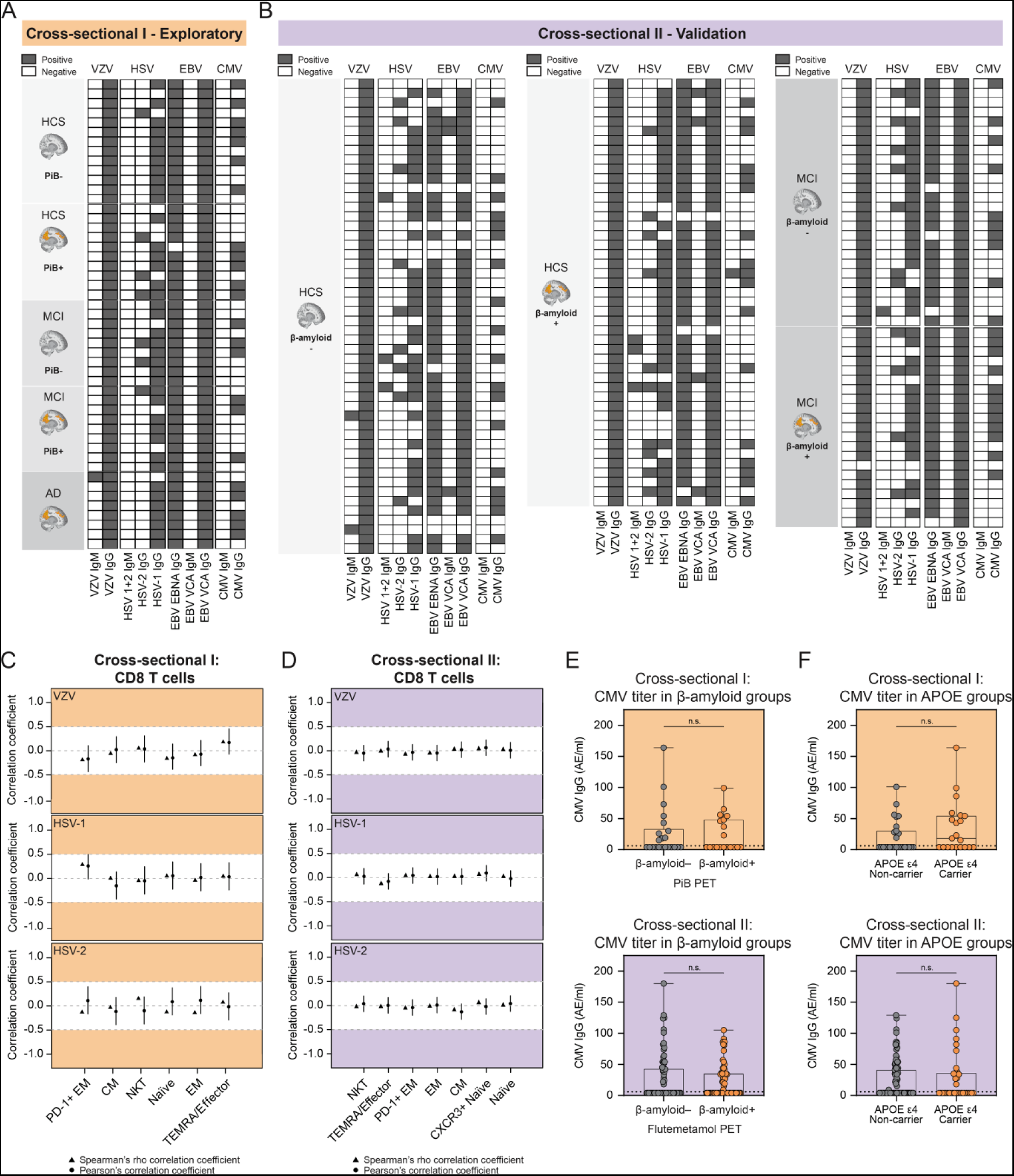
Herpesvirus Infection Status in Cross-sectional Cohorts. **(A+B)** Qualitative status of antiviral antibody titers against (re-)activated *Herpesviridae* for all individual subjects of *Cross-sectional I* **(A)** and *Cross-sectional II* **(B)**. Antibody titers are subdivided into IgM titers (early phase antibodies, acute infection) and IgG titers (long-term immunity, higher levels after recent infections) against Varicella-Zoster-Virus (VZV), Herpes simplex virus (HSV) 1+2, Epstein-Barr nuclear *antigen* (EBNA) of Epstein-Barr virus (EBV), viral capsid antigen (VCA) of EBV and Cytomegalovirus (CMV). **(C+D)** Correlations between herpesvirus IgG titers and relative CD8^+^ T cell cluster abundances for *Cross-sectional I* **(C)** and *Cross-sectional II* **(D)**. Correlations for VZV, HSV-1 and HSV-2 IgG titers are displayed. No substantial differences have been found. **(E)** Quantification of IgG titers against CMV (in AE/ml, threshold ≥6) in PET β-amyloid+ and β-amyloid– subjects of *Cross-sectional I* and *Cross-sectional II*. No differences have been found. **(F)** Quantification of IgG titers against CMV (in AE/ml, threshold ≥6) in APOE ε4 carriers and non-carriers of *Cross-sectional I* and *Cross-sectional II*. No differences have been found. Black dots represent the estimated Pearson’s r correlation coefficients and the black lines the corresponding 95% confidence intervals. Black triangles represent the estimated Spearman’s rho correlation coefficients. Dashed lines are indicating a correlation cut-off at rho = 0.5 or rho = -0.5.

**Figure S5.**
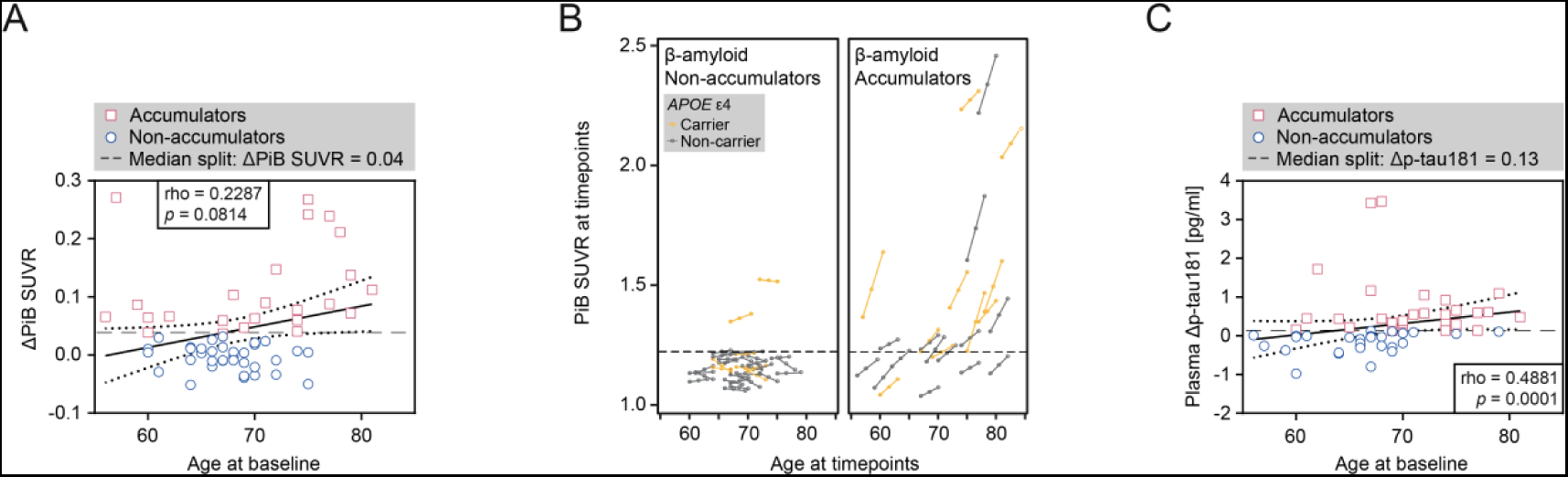
Longitudinal Study: AD Biomarkers versus Age. **(A)** Correlation of ΔPiB SUVR with individual age at baseline with linear regression; horizontal dashed line represents cut-off for β-amyloid Accumulators (ΔPiB SUVR ≥ 0.04). **(B)** Plot of individual PiB SUVR values versus corresponding ages at all three time points for β-amyloid Non-accumulators and Accumulators. PiB SUVR values at *Follow-up1* time point are individually imputed based on linear interpolation. Dashed line indicates cut-off for cerebral β-amyloid-positive subjects (PiB SUVR = 1.223). Orange symbols mark subjects carrying at least one *APOE* ε4 allele. **(C)** Correlation of plasma Δp-tau181 with individual age at baseline with linear regression; horizontal dashed line represents cut-off for p-tau181 Accumulators (Δp-tau181 ≥ 0.13). Each symbol in quantification graphs represents data from one subject. All *p* values ≤ 0.1 (FDR 10%) are displayed. Correlation coefficients (S5A, S5C) were calculated via Spearman approach (rho and *p* value reported). Dotted lines represent 95% confidence intervals of a linear regression.

**Figure S6.**
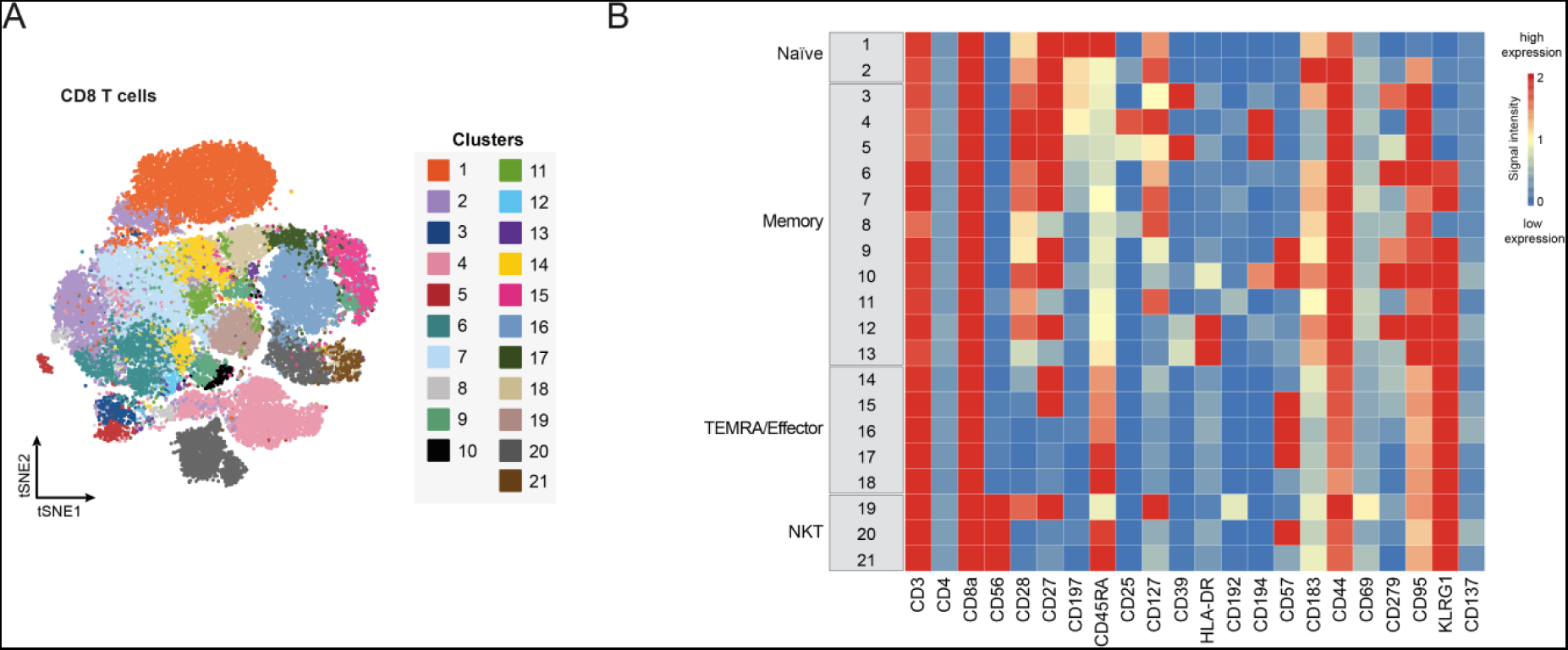
Longitudinal Study: tSNE Plot and Heatmap for CD8^+^ T cell Populations. **(A)** tSNE map and FlowSOM clustering for all PBMC-derived CD8^+^ T cells averaged over all subject samples of the longitudinal study (tSNE settings: iterations = 4’000, events = max. 10’000, perplexity = 50; FlowSOM settings: k25, merged into 21 subclusters). **(B)** Heatmap for identification of 21 CD8^+^ T cell clusters. Categories of CD8^+^ T cell clusters are allocated according to similar marker expression within major CD8^+^ T cell subpopulations found in the blood. Heatmap shows arcsinh-transformed median marker intensities.

## Notes

https://zenodo.org/record/7157682

