## Supplemental Tables for "Early β-amyloid accumulation in the brain is associated with peripheral T cell alterations"

**Table S1. Detailed group sizes, demographics and characterization of the cross-sectionally analyzed exploratory cohort (*Cross-sectional I*)**

| | **Total** | **HCS β-amyloid–** | **HCS β-amyloid+** | **MCI β-amyloid–** | **MCI β-amyloid+** | **AD** | ***P* value for statistical test** |
|---|---|---|---|---|---|---|---|
| **Subjects** | 50 | 13 | 10 | 9 | 9 | 9 | |
| **Age** | 75.1 (6.5) | 73.4 (3.2) | 76.3 (3.7) | 70.0 (7.4) | 74.1 (5.3) | 82.3 (6.8) **$p < 0.01$** | **$P < 0.001$** |
| **Years of education** | 14.4 (3.1) | 15.2 (3.4) | 14.0 (2.7) | 15.9 (2.9) | 14.7 (3.4) | 12.2 (2.3) | $P = 0.1107$ |
| **Sex female / male** | 18 / 32 | 4 / 9 | 4 / 6 | 3 / 6 | 2 / 7 | 5 / 4 | $P = 0.6470$ |
| ***APOE* ε4 carrier vs. non-carrier** | 22 / 27 | 3 / 10 | 5 / 4 (one case unknown) | 3 / 6 | 6 / 3 | 5 / 4 | $P = 0.2363$ |
| **MMSE** | 27.08 (4.7) | 29.46 (0.5) | 29.30 (1.3) | 28.89 (1.2) | 27.22 (2.4) | 18.78 (5.6) **$p < 0.0001$** | **$P < 0.0001$** |
| **CDR** | 0.33 (0.5) | 0 (0) | 0 (0) | 0.17 (0.3) | 0.39 (0.2) **$p < 0.05$** | 1.28 (0.6) **$p < 0.0001$** | **$P < 0.0001$** |
| **PiB SUVR** | 1.22 [0.73] | 1.15 [0.09] | 1.64 [0.47] **$p < 0.001$** | 1.15 [0.01] | 2.19 [0.47] **$p < 0.0001$** | - | **$P < 0.0001$** |
| **PiB Centiloid** | 5.46 [69.44] | -0.99 [5.82] | 59.01 [30.13] **$p < 0.001$** | 0.20 [7.32] | 95.54 [27.86] **$p < 0.0001$** | - | **$P < 0.0001$** |
| **Plasma Aβ42/Aβ40 ratio** | 0.047 [0.011] | 0.055 [0.002] | 0.044 [0.005] **$p < 0.01$** | 0.051 [0.009] | 0.043 [0.006] **$p < 0.0001$** | 0.044 [0.005] **$p < 0.05$** | **$P < 0.001$** |
| **Plasma total tau in pg/ml** | 2.20 [1.25] | 1.69 [0.63] | 2.41 [1.32] **$p < 0.05$** | 2.46 [0.85] | 1.76 [0.65] | 3.29 [0.36] **$p < 0.001$** | **$P < 0.01$** |
| **Plasma p-tau181 in pg/ml** | 1.89 [1.23] | 1.42 [0.43] | 2.24 [0.46] **$p < 0.05$** | 1.21 [0.56] | 2.52 [1.13] **$p < 0.01$** | 3.10 [1.65] **$p < 0.01$** | **$P < 0.001$** |

Age (mean ± standard deviation (SD)), years of education (mean ± SD), self-reported gender, apolipoprotein E (*APOE*) ε4 gene distribution, mini-mental state examination (MMSE) score (mean ± SD), clinical dementia rating (CDR) (global score, mean ± SD), cortical PiB SUVR (median with [IQR]), cortical PiB centiloid (median with [IQR]) and plasma levels of Aβ42/Aβ40, total tau, and p-tau181 (median with [IQR]) for each group are reported. The overall *P* values of the applied statistical tests are shown separately. The individual, adjusted *p* values indicate significant difference from HCS PiB- group and were calculated using parametric one-way ANOVA with Bonferroni's multiple comparison correction (for parameters 'Age', 'Years of education', 'MMSE' and 'CDR') or non-parametric Kruskal-Wallis test with false-discovery-rate (FDR) method of Benjamini and Hochberg (for parameters 'PiB SUVR', 'PiB Centiloid' and plasma biomarkers). The *P* values for categorical variables such as 'Sex' and '*APOE* ε4 carrier' were calculated via Chi-square test for independence. Significant values are displayed in bold.

**Table S2. Detailed group sizes, demographics and characterization of the cross-sectionally analyzed validation cohort (*Cross-sectional II*)**

| | Total | HCS β-amyloid– | HCS β-amyloid+ | MCI β-amyloid– | MCI β-amyloid+ | *P* value for statistical test |
|---|---|---|---|---|---|---|
| **Subjects** | 142 | 50 | 45 | 26 | 21 | |
| **Age** | 68.5 (7.5) | 66.5 (6.3) | 67.3 (6.7) | 70.1 (9.1) | 73.9 (7.1) ***p* < 0.001** | ***P* < 0.001** |
| **Years of education** | 15.29 | 15.52 (2.9) | 16.33 (2.8) | 14.54 (3.0) | 14.76 (3.2) | *P* = 0.0564 |
| **Sex female / male** | 46 / 86 | 27 / 23 | 10 / 35 | 9 / 17 | 10 / 11 | ***P* < 0.05** |
| ***APOE* ε4 carrier vs. non-carrier** | 35 / 107 | 10 / 40 | 17 / 28 | 1 / 25 | 7 / 14 | ***P* < 0.01** |
| **MMSE** | 28.74 | 29.48 (0.8) | 29.07 (1.3) | 28.54 (1.1) ***p* < 0.01** | 27.86 (1.7) ***p* < 0.0001** | ***P* < 0.0001** |
| **CDR** | 0.13 | 0.03 (0.1) | 0.02 (0.1) | 0.15 (0.2) ***p* < 0.01** | 0.33 (0.2) ***p* < 0.0001** | ***P* < 0.0001** |
| **FMM SUVR** | 1.24 [0.21] | 1.15 [0.10] | 1.36 [0.17] ***p* < 0.0001** | 1.16 [0.11] | 1.54 [0.64] ***p* < 0.0001** | ***P* < 0.0001** |
| **FMM Centiloid** | 8.40 [20.01] | 1.24 [8.57] | 19.98 [18.34] ***p* < 0.0001** | 0.67 [11.16] | 38.92 [58.49] ***p* < 0.0001** | ***P* < 0.0001** |

Age (mean ± standard deviation (SD)), years of education (mean ± SD), self-reported gender, apolipoprotein E (*APOE*) ε4 distribution, mini-mental state examination (MMSE) score (mean ± SD), clinical dementia rating (CDR) (global score, mean ± SD), cortical Flutemetamol (FMM) SUVR (median with [IQR]) and cortical FMM centiloid (median with [IQR]) for each group are reported. The overall *P* values of the applied statistical tests are shown separately. The individual, adjusted *p* values indicate significant difference from HCS β-amyloid– group and were calculated using parametric one-way ANOVA with Bonferroni's multiple comparison correction (for parameters 'Age', 'Years of education', 'MMSE' and 'CDR') or non-parametric Kruskal-Wallis test with false-discovery-rate (FDR) method of Benjamini and Hochberg (for parameters 'FMM SUVR' and 'FMM Centiloid'). The *P* values for categorical variables such as 'Sex' and '*APOE* ε4 carrier' were calculated via Chi-square test for independence. Significant values are displayed in bold.

**Table S3. Longitudinal study - PBMC sample availability at different time points**

| Subject Identifier | β-amyloid Group | Baseline | Follow-up 1 | Follow-up 2 |
|---|---|---|---|---|
| LS_1 | Accumulator | Yes | Yes | Yes |
| LS_2 | Accumulator | No | No | Yes |
| LS_3 | Accumulator | No | Yes | Yes |
| LS_4 | Non-accumulator | Yes | No | Yes |
| LS_5 | Accumulator | Yes | Yes | Yes |
| LS_6 | Accumulator | Yes | No | Yes |
| LS_7 | Non-accumulator | Yes | Yes | Yes |
| LS_8 | Non-accumulator | Yes | No | Yes |
| LS_9 | Non-accumulator | Yes | Yes | Yes |
| LS_10 | Non-accumulator | Yes | No | Yes |
| LS_11 | Non-accumulator | Yes | Yes | Yes |
| LS_12 | Accumulator | Yes | No | No |
| LS_13 | Accumulator | No | No | Yes |
| LS_14 | Non-accumulator | Yes | No | Yes |
| LS_15 | Non-accumulator | Yes | Yes | No |
| LS_16 | Non-accumulator | Yes | Yes | Yes |
| LS_17 | Non-accumulator | Yes | No | Yes |
| LS_18 | Accumulator | No | Yes | Yes |
| LS_19 | Accumulator | No | Yes | Yes |
| LS_20 | Non-accumulator | No | Yes | Yes |
| LS_21 | Non-accumulator | No | Yes | Yes |
| LS_22 | Accumulator | No | Yes | Yes |
| LS_23 | Non-accumulator | No | No | Yes |
| LS_24 | Non-accumulator | No | Yes | Yes |
| LS_25 | Non-accumulator | Yes | Yes | Yes |
| LS_26 | Non-accumulator | No | Yes | Yes |
| LS_27 | Non-accumulator | No | Yes | Yes |
| LS_28 | Non-accumulator | No | Yes | Yes |
| LS_29 | Non-accumulator | No | Yes | Yes |
| LS_30 | Non-accumulator | No | Yes | Yes |
| LS_31 | Non-accumulator | No | Yes | Yes |
| LS_32 | Non-accumulator | No | Yes | Yes |
| LS_33 | Accumulator | No | Yes | Yes |
| LS_34 | Accumulator | Yes | Yes | Yes |
| LS_35 | Non-accumulator | No | Yes | Yes |
| LS_36 | Non-accumulator | No | Yes | Yes |
| LS_37 | Accumulator | No | Yes | Yes |
| LS_38 | Non-accumulator | Yes | Yes | Yes |
| LS_39 | Accumulator | Yes | Yes | Yes |

| | | | | |
|---|---|---|---|---|
| LS_40 | Non-accumulator | Yes | Yes | Yes |
| LS_41 | Non-accumulator | No | Yes | Yes |
| LS_42 | Accumulator | No | Yes | Yes |
| LS_43 | Accumulator | No | Yes | Yes |
| LS_44 | Accumulator | No | Yes | Yes |
| LS_45 | Non-accumulator | No | Yes | Yes |
| LS_46 | Accumulator | Yes | Yes | Yes |
| LS_47 | Non-accumulator | Yes | Yes | Yes |
| LS_48 | Non-accumulator | Yes | Yes | Yes |
| LS_49 | Accumulator | No | Yes | Yes |
| LS_50 | Accumulator | No | Yes | Yes |
| LS_51 | Accumulator | No | Yes | Yes |
| LS_52 | Non-accumulator | No | Yes | Yes |
| LS_53 | Non-accumulator | No | Yes | Yes |
| LS_54 | Non-accumulator | Yes | Yes | Yes |
| LS_55 | Non-accumulator | No | Yes | Yes |
| LS_56 | Accumulator | Yes | Yes | Yes |
| LS_57 | Accumulator | Yes | Yes | Yes |
| LS_58 | Non-accumulator | Yes | No | Yes |
| LS_59 | Accumulator | No | Yes | Yes |

**Table S4. Detailed group sizes, demographics and characterization of the longitudinally analyzed cohort**

| | Total | Cerebral β-amyloid Non-accumulators | Cerebral β-amyloid Accumulators | *P* value |
|---|---|---|---|---|
| **Subjects** | 59 | 35 | 24 | |
| **Age at Baseline** | 68.47 (5.7) | 67.40 (3.9) | 70.04 (7.6) | *P* = 0.0840 |
| **Years of education** | 15.03 (2.8) | 15.11 (3.0) | 14.92 (2.5) | *P* = 0.7995 |
| **Sex female / male** | 27 / 32 | 14 / 21 | 13 / 11 | *P* = 0.3027 |
| ***APOE* ε4 carrier vs. non-carrier** | 18 / 41 | 8 / 27 | 10 / 14 | *P* = 0.1555 |
| **MMSE Baseline** | 29.25 (1.0) | 29.29 (1.0) | 29.21 (0.9) | *P* = 0.7546 |
| **MMSE Follow-up1** | 29.41 (0.7) | 29.50 (0.6) | 29.29 (0.8) | *P* = 0.2541 |
| **MMSE Follow-up2** | 29.22 (0.8) | 29.26 (0.8) | 29.17 (0.8) | *P* = 0.6728 |
| **PiB SUVR Baseline** | 1.18 [0.09] | 1.16 [0.07] | 1.23 [0.22] | ***P* < 0.01** |
| **PiB SUVR Follow-up2** | 1.19 [0.15] | 1.15 [0.09] | 1.30 [0.34] | ***P* < 0.0001** |
| **ΔPiB SUVR** | 0.02 [0.07] | 0 [0.03] | 0.08 [0.08] | ***P* < 0.0001** |
| **Plasma p-tau181 in pg/ml Baseline** | 1.35 [0.59] | 1.34 [0.55] | 1.49 [0.84] | *P* = 0.1588 |
| **Plasma p-tau181 in pg/ml Follow-up2** | 1.39 [0.95] | 1.30 [0.72] | 1.74 [1.46] | ***P* < 0.05** |
| **Plasma Δp-tau181 in pg/ml** | 0.13 [0.53] | 0 [0.59] | 0.23 [0.48] | ***P* < 0.05** |

Age at *Baseline* (mean ± SD), years of education (mean ± SD), self-reported gender, *APOE* ε4 distribution, MMSE score at all time points (mean ± SD), cortical PiB SUVR and plasma p-tau181 at *Baseline* and *Follow-up2* time points (median with [IQR]), as well as ΔPiB SUVR and Δp-tau181 (*Follow-up2 – Baseline*, median with [IQR]) for each group are reported. Bold *P* values indicate significant difference from Non-accumulators group and were calculated using parametric *t* test (for parameters 'Age at *Baseline*', 'Years of education' and 'MMSE *Baseline/Follow-up1/Follow-up2*') or nonparametric Mann-Whitney test (for parameters 'PiB SUVR *Baseline*', 'PiB SUVR *Follow-up2*' and 'ΔPiB SUVR', as well as 'p-tau181 *Baseline*', 'p-tau181 *Follow-up2*' and 'Δp-tau181'). The *P* values for categorical variables such as 'Sex' and '*APOE* ε4 carrier' were calculated via Fisher's exact test to prove independence.

**Table S5. Neuropsychological tests**

| Predefined order | Test name | Modifications | Source reference |
|---|---|---|---|
| 1 | CERAD Battery, German version | Addition of 'Victoria Stroop test' and 'Kramer card sorting test' | **(Thalmann et al., 2000)** (CERAD) **(Stroop, 1935)** (Stroop test) **(Kramer, 1954)** (Card sorting test) |
| 2 [a] | Phonemic Fluency over 3 min (Letter: S) | - | **(Delis et al., 2001)** |
| 3 | Visual Pair Learning of the WMS-R Battery | - | **(Härting et al., 2000)** |
| 4 [a] | RAVLT (VLMT), German version | - | **(Helmstaedter et al., 2001)** |
| 5 | Rey-Osterrieth Complex Figure Test | With incidental immediate recall and 30' late recall; Taylor criteria used for evaluation | **(Osterrieth, 1944)** **(Taylor, 1969)** |
| 6 [a] | Digit Span and Corsi Block-Tapping Task | Forward and backward * tasks | **(Corsi, 1972)** **(Härting et al., 2000)** |
| 7 [a] | Non-verbal Fluency (5-point Test) | - | **(Regard et al., 1982)** |
| 8 | Simple Two-dimensional Mental Rotation Task | - | **(Shepard and Metzler, 1971)** |
| 9 | Trail Making Test | Part A and B | **(Reitan, 1955)** |
| 10 | Clock Test | - | **(Manos and Wu, 1994)** |

[a] used for the final analysis of cognitive functions in the cognitively healthy cohort only

**Table S6. Live cell barcoding panels**

| *Cross-sectional I* | | | |
|---|---|---|---|
| **Metal Tag** | **Target** | **Clone** | **Source** |
| $^{89}Y$ | CD45 | HI30 | Fluidigm |
| $^{143}Nd$ | CD45 | HI30 | In-house labeling |
| $^{148}Nd$ | CD45 | HI30 | In-house labeling |
| $^{152}Sm$ | CD45 | HI30 | In-house labeling |
| $^{154}Sm$ | CD45 | HI30 | Fluidigm |
| $^{162}Dy$ | CD45 | HI30 | In-house labeling |
| $^{168}Er$ | CD45 | HI30 | In-house labeling |
| $^{173}Yb$ | CD45 | HI30 | In-house labeling |
| *Cross-sectional II* | | | |
| $^{89}Y$ | CD45 | HI30 | Fluidigm |
| $^{106}Cd$ | CD45 | HI30 | In-house labeling |
| $^{110}Cd$ | CD45 | HI30 | In-house labeling |
| $^{111}Cd$ | CD45 | HI30 | In-house labeling |
| $^{112}Cd$ | CD45 | HI30 | In-house labeling |
| $^{113}In$ | CD45 | HI30 | In-house labeling |
| $^{114}Cd$ | CD45 | HI30 | In-house labeling |
| $^{169}Tm$ | CD45 | HI30 | In-house labeling |
| $^{171}Yb$ | CD45 | HI30 | In-house labeling |
| $^{173}Yb$ | CD45 | HI30 | In-house labeling |
| **Longitudinal study** | | | |
| $^{89}Y$ | CD45 | HI30 | Fluidigm |
| $^{113}In$ | CD45 | HI30 | In-house labeling |
| $^{115}In$ | CD45 | HI30 | In-house labeling |
| $^{162}Dy$ | CD45 | HI30 | In-house labeling |
| $^{163}Dy$ | CD45 | HI30 | In-house labeling |
| $^{168}Er$ | CD45 | HI30 | In-house labeling |
| $^{169}Tm$ | CD45 | HI30 | In-house labeling |
| $^{171}Yb$ | CD45 | HI30 | In-house labeling |
| $^{173}Yb$ | CD45 | HI30 | In-house labeling |
| $^{175}Lu$ | CD45 | HI30 | In-house labeling |

Live-cell barcoding panel for T cell study in *Cross-sectional I* included 8 anti-CD45 antibodies of the same clone coupled with 8 different heavy metal isotopes. We selected combinations of 3 antibodies for barcoding (= 8-choose-3 approach, 56 possible combinations). Live-cell barcoding for *Cross-sectional II* and the longitudinal T and B cell studies included 10 anti-CD45 antibodies, allowing for a 10-choose-4 approach (210 possible combinations).
