## Supplemental Materials for "Early β-amyloid accumulation in the brain is associated with peripheral T cell alterations"

| REAGENT or RESOURCE | SOURCE | IDENTIFIER |
| --- | --- | --- |
| Antibodies - Cross-sectional study I | | |
| Anti-Human CD19 (HIB19)-142Nd Antibody | Fluidigm | Cat# 3142001, RRID: AB_2651155 |
| Anti-Human BDCA1/CD1c (AD5-8E7) Purified Antibody; In-House Labeling with 144Nd | Miltenyi Biotec | Cat# 130-090-695, RRID: N/A |
| Anti-Human CD4 (RPA-T4)-145Nd Antibody | Fluidigm | Cat# 3145001, RRID: AB_2661789 |
| Anti-Human CD8a (IA6-2)-146Nd Antibody | Fluidigm | Cat# 3146003B, RRID: AB_2687833 |
| Anti-Human CD303/BDCA2 (201A)-147Sm Antibody | Fluidigm | Cat# 3147009B, RRID: N/A |
| Anti-Human CD25/IL-2R (2A3)-149Sm Antibody | Fluidigm | Cat# 3149010B, RRID: AB_2756416 |
| Anti-Human CD141/BDCA3 (AD5-14H12) Purified Antibody; In-House Labeling with 150Nd | Miltenyi Biotec | Cat# 130-090-694, RRID: N/A |
| Anti-Human CD123/IL-3R (6H6)-151Eu Antibody | Fluidigm | Cat# 3151001, RRID: AB_2661794 |
| Anti-Human CD192/CCR2 (K036C2)-153Eu Antibody | Fluidigm | Cat# 3153023B, RRID: N/A |
| Anti-Human CD279/PD-1 (EH12.2H7)-155Gd Antibody | Fluidigm | Cat# 3155009B, RRID: AB_2687854 |
| Anti-Human CD183/CXCR3 (G025H7)-156Gd Antibody | Fluidigm | Cat# 3156004B, RRID: N/A |
| Anti-Human CD194/CCR4 (L291H4)-158Gd Antibody | Fluidigm | Cat# 3158032A, RRID: AB_2893003 |
| Anti-Human CD197/CCR7 (G043H7)-159Tb Antibody | Fluidigm | Cat# 3159003, RRID: AB_2714155 |
| Anti-Human CD28 (CD28.2)-160Gd Antibody | Fluidigm | Cat# 3160003B, RRID: N/A |
| Anti-Human CD127/IL-7Ra (A019D5) Purified Antibody; In-House Labeling with 164Dy | Biolegend | Cat# 351337, RRID: N/A |
| Anti-Human CD39 (A1) Purified Antibody; In-House Labeling with 165Ho | Biolegend | Cat# 328221, RRID: N/A |
| Anti-Human CD44 (BJ18)-166Er Antibody | Fluidigm | Cat# 3166001B, RRID: AB_2744692 |
| Anti-Human CD27 (O323)-167Er Antibody | Fluidigm | Cat# 3167002B, RRID: N/A |
| Anti-Human CD45RA (HI100)-169Tm Antibody | Fluidigm | Cat# 3169008B, RRID: N/A |
| Anti-Human CD3 (UCHT1)-170Er Antibody | Fluidigm | Cat# 3170001, RRID: AB_2661807 |
| Anti-Human CD69 (FN50) Purified Antibody; In-House Labeling with 171Yb | Biolegend | Cat# 310902, RRID: N/A |
| Anti-Human CD57 (HCD57)-172Yb Antibody | Fluidigm | Cat# 3172009B, RRID: N/A |
| Anti-Human HLA-DR (L243)-174Yb Antibody | Fluidigm | Cat# 3174001, RRID: AB_2665397 |
| Anti-Human CD14 (M5E2)-175Lu Antibody | Fluidigm | Cat# 3175015B, RRID: AB_2811083 |
| Anti-Human CD56 (NCAM16.2)-176Yb Antibody | Fluidigm | Cat# 3176008, RRID: AB_2661813 |

| | | |
|---|---|---|
| Anti-Human CD16 (3G8)-209Bi Antibody | Fluidigm | Cat# 3209002B, RRID: AB_2756431 |
| Antibodies - Cross-sectional study II and Longitudinal study | | |
| Anti-Human CD45RA (HI100) Purified Antibody; In-House labeling with 141Pr | Biolegend | Cat# 304102, RRID: N/A |
| Anti-Human CD19 (HIB19)-142Nd Antibody | Fluidigm | Cat# 3142001, RRID: AB_2651155 |
| Anti-Human CD123/IL-3R (6H6)-143Nd Antibody | Fluidigm | Cat# 3143014B, RRID: AB_2811081 |
| Anti-Human CD69 (FN50)-144Nd Antibody | Fluidigm | Cat# 3144018, RRID: AB_2687849 |
| Anti-Human CD4 (RPA-T4)-145Nd Antibody | Fluidigm | Cat# 3145001, RRID: AB_2661789 |
| Anti-Human CD8a (IA6-2)-146Nd Antibody | Fluidigm | Cat# 3146003B, RRID: AB_2687833 |
| Anti-Human CD11a (HI111) Purified Antibody; In-House labeling with 146Nd | Biolegend | Cat# 301202, RRID: N/A |
| Anti-Human CD303/BDCA2 (201A)-147Sm Antibody | Fluidigm | Cat# 3147009B, RRID: N/A |
| Anti-Human CD137/4-1BB (4B4-1) Purified Antibody; In-House labeling with 148Nd | Biolegend | Cat# 309802, RRID: N/A |
| Anti-Human CD25/IL-2R (2A3)-149Sm Antibody | Fluidigm | Cat# 3149010B, RRID: AB_2756416 |
| Anti-Human CD127/IL-7Ra (A019D5) Purified Antibody; In-House Labeling with 150Nd | Biolegend | Cat# 351302, RRID: N/A |
| Anti-Human CD14 (M5E2)-151Eu Antibody | Fluidigm | Cat# 3151009B, RRID: AB_2810244 |
| Anti-Human KLRG1 (13F12F2) Purified Antibody; In-House labeling with 152Sm | ThermoFisher | Cat# 16-9488-85, RRID: N/A |
| Anti-Human CD192/CCR2 (K036C2)-153Eu Antibody | Fluidigm | Cat# 3153023B, RRID: N/A |
| Anti-Human BDCA1/CD1c (AD5-8E7) Purified Antibody; In-House Labeling with 154Sm | Miltenyi Biotec | Cat# 130-090-695, RRID: N/A |
| Anti-Human CD279/PD-1 (EH12.2H7)-155Gd Antibody | Fluidigm | Cat# 3155009B, RRID: AB_2687854 |
| Anti-Human CD183/CXCR3 (G025H7)-156Gd Antibody | Fluidigm | Cat# 3156004B, RRID: N/A |
| Anti-Human CD194/CCR4 (L291H4)-158Gd Antibody | Fluidigm | Cat# 3158032A, RRID: AB_2893003 |
| Anti-Human CD197/CCR7 (G043H7)-159Tb Antibody | Fluidigm | Cat# 3159003, RRID: AB_2714155 |
| Anti-Human CD28 (CD28.2)-160Gd Antibody | Fluidigm | Cat# 3160003B, RRID: AB_2868400 |
| Anti-Human CD152 (14D3)-161Dy Antibody | Fluidigm | Cat# 3161004B, RRID: N/A |
| Anti-Human TCRγ/δ (B1) Purified Antibody; In-House Labeling with 162Dy | Biolegend | Cat# 331202, RRID: N/A |
| Anti-Human CD95 (DX2)-164Dy Antibody | Fluidigm | Cat# 3164008B, RRID: N/A |
| Anti-Human CD39 (A1) Purified Antibody; In-House Labeling with 165Ho | Biolegend | Cat# 328221, RRID: N/A |
| Anti-Human CD44 (BJ18)-166Er Antibody | Fluidigm | Cat# 3166001B, RRID: AB_2744692 |
| Anti-Human CD27 (L128)-167Er Antibody | Fluidigm | Cat# 3167006B, |

| | | |
|---|---|---|
| | | RRID: N/A |
| Anti-Human CD73 (AD2)-168Er Antibody | Fluidigm | Cat# 3168015B, RRID: AB_2810249 |
| Anti-Human CD3 (UCHT1)-170Er Antibody | Fluidigm | Cat# 3170001, RRID: AB_2661807 |
| Anti-Human CD57 (HCD57)-172Yb Antibody | Fluidigm | Cat# 3172009B, RRID: AB_2888930 |
| Anti-Human HLA-DR (L243)-174Yb Antibody | Fluidigm | Cat# 3174001, RRID: AB_2665397 |
| Anti-Human CD278/ICOS (C398.4A)-175Lu Antibody | Fluidigm | Cat# 3175039B, RRID: AB_2905647 |
| Anti-Human CD56 (NCAM16.2)-176Yb Antibody | Fluidigm | Cat# 3176008, RRID: AB_2661813 |
| Anti-Human CD16 (3G8)-209Bi Antibody | Fluidigm | Cat# 3209002B, RRID: AB_2756431 |
| Chemicals, Peptides, and Recombinant Proteins | | |
| EQ™ Four Element Calibration Beads | Fluidigm | Cat# 201078 |
| Tuning Solution | Fluidigm | Cat# 201072 |
| Maxpar® Water | Fluidigm | Cat# 201069 |
| Maxpar® Cell Staining Buffer | Fluidigm | Cat# 201068 |
| Maxpar® Fix and Perm Buffer | Fluidigm | Cat# 201067 |
| Maxpar® Cell Acquisition Solution (CAS) | Fluidigm | Cat# 201241 |
| Cell-ID™ Cisplatin-$^{198}$Pt | Fluidigm | Cat# 201064 |
| Cell-ID™ Intercalator-Ir | Fluidigm | Cat# 201192 |
| Human TruStain FcX™ (Fc Receptor Blocking Solution) | Biolegend | Cat# 422302 |
| RPMI-1640 Cell Culture Medium | Sigma Aldrich | Cat# R0883 |
| Glutamax Supplement | Gibco/Thermo Fisher | Cat# 35050061 |
| Fetal Bovine Serum | Gibco/Thermo Fisher | Cat# 10500064 |
| Dimethyl Sulfoxide (DMSO) | Thermo Fisher | Cat# D12345 |
| Pierce™ 16% Formaldehyde (w/v) | Thermo Fisher | Cat# 28908 |
| Critical Commercial Assays | | |
| Maxpar® X8 Multimetal Labeling Kit | Fluidigm | Cat# 201300 |
| Cell-ID™ 20-Plex Pd Barcoding Kit | Fluidigm | Cat# 201060 |
| Deposited Data | | |
| Datasets and code for LMM | Zenodo | http://doi.org/10.5281/zenodo.4911274 |
| Software and Algorithms | | |
| Graphpad Prism (Version 8.4.2) | GraphPad Software | https://www.graphpad.com/scientific-software/prism/ |
| Cytobank Premium (Version 7.3.0) | Beckman Coulter | https://premium.cytobank.org/cytobank/ |
| R (Version 4.0.2) | R Core Team | https://cran.r-project.org/src/base/R-4/ |
| RStudio (Version 1.3.1093) | RStudio, Inc. | https://rstudio.com/ |
| flowCore (Version 2.0.1) | (Hahne et al., 2009) (Ellis et al., 2009) | https://bioconductor.org/packages/release/bioc/html/flowCore.html |

| | | |
|---|---|---|
| ggplot2 (Version 3.3.2) | (Wickham et al., 2016) | https://github.com/tidyverse/ggplot2/releases |
| Rtsne (Version 0.15) | (Van der Maaten and Hinton, 2008) | https://github.com/jkrijthe/Rtsne |
| FlowSOM (Version 1.20.0) | (Van Gassen et al., 2015) | http://bioconductor.org/packages/release/bioc/html/FlowSOM.html |
| dplyr (Version 1.0.2) | (Wickham et al., 2015) | https://CRAN.R-project.org/package=dplyr |
| Premessa (Version 0.2.6) | (Gherardini, 2019) | https://github.com/ParkerICI/premessa |
| Catalyst (Version 1.14.0) | (Crowell et al., 2020) | https://bioconductor.org/packages/release/bioc/html/CATALYST.html<br><br>https://github.com/HelenaLC/CATALYST |
| lme4 (Version 1.1-26) | (Bates et al., 2015) | https://github.com/lme4/lme4/ |
| Other | | |
| Helios CyTOF2 mass cytometer | DVS Sciences, Fluidigm | N/A |
