## Supplemental Formula S1 for "Early β-amyloid accumulation in the brain is associated with peripheral T cell alterations"

**Formula S1: Linear Mixed Model (LMM)**

$$\boldsymbol{y} = \boldsymbol{X}\boldsymbol{\beta} + \boldsymbol{Z}\boldsymbol{u} + \boldsymbol{\varepsilon} \tag{1}$$

**fixed effects**

- $n$: number of observations per cluster (for each subject up to two time points).
- $\boldsymbol{y} \in \mathbb{R}^{n\times 1}$: response vector (cortical PiB SUVR or plasma p-tau181 concentration of each timepoint considered).
- $\boldsymbol{X} \in \mathbb{R}^{n\times 5}$: design matrix for fixed effects, i.e. matrix where a column corresponds to the Intercept or a covariate (Age; Relative Abundance of immune cells at each timepoint considered; Group (Accumulator vs. Non-accumulator); Interaction between Relative Abundance and Group (Accumulator vs. Non-accumulator)).
- $\boldsymbol{\beta} \in \mathbb{R}^{5\times 1}$: column vector of fixed effect regression coefficients.

**random effects**

- $J \in \mathbb{N}$: groups for the random effects (J: number of individual subjects).
- $\boldsymbol{Z} \in \mathbb{R}^{n\times J}$: design matrix for random effects (random intercept per subject).
- $\boldsymbol{u} \in \mathbb{R}^{J\times 1}$: column vector of random effect coefficients.

**residual**

- $\boldsymbol{\varepsilon} \in \mathbb{R}^{n\times 1}$: column vector of residuals, the part of $\boldsymbol{y}$ which is not explained by $\boldsymbol{X}\boldsymbol{\beta} + \boldsymbol{Z}\boldsymbol{u}$

**see** (Singer & Willett (2003).
